# Seasonal Shifts in Community Composition and Proteome Expression in a Sulfur-Cycling Cyanobacterial Mat

**DOI:** 10.1101/2023.01.30.526236

**Authors:** Sharon L Grim, Dack G Stuart, Phoebe Aron, Naomi E Levin, Lauren E Kinsman-Costello, Jacob E Waldbauer, Gregory J Dick

## Abstract

Seasonal changes in light and physicochemical conditions have strong impacts on cyanobacteria, but how they affect community structure, metabolism, and biogeochemistry of cyanobacterial mats remains unclear. Light may be particularly influential for cyanobacterial mats exposed to sulfide by altering the balance of oxygenic photosynthesis and sulfide-driven anoxygenic photosynthesis. We studied temporal shifts in irradiance, water chemistry, and community structure and function of microbial mats in Middle Island Sinkhole (MIS), where anoxic and sulfate-rich groundwater provides habitat for cyanobacteria that conduct both oxygenic and anoxygenic photosynthesis. Seasonal changes in light and groundwater chemistry were accompanied by shifts in bacterial community composition, with a succession of dominant cyanobacteria from *Phormidium* to *Planktothrix,* and an increase in diatoms, sulfur-oxidizing bacteria, and sulfate-reducing bacteria from summer to autumn. Differential abundance of cyanobacterial light harvesting proteins likely reflects a physiological response of cyanobacteria to light level. *Beggiatoa* sulfur oxidation proteins were more abundant in autumn. Correlated abundances of taxa through time suggest interactions between sulfur oxidizers and sulfate reducers, sulfate reducers and heterotrophs, and cyanobacteria and heterotrophs. These results support the conclusion that seasonal change, including light availability, has a strong influence on community composition and biogeochemical cycling of sulfur and O_2_ in cyanobacterial mats.

**Originality-Significance Statement:** Cyanobacterial mats are found in terrestrial and aquatic environments on modern Earth and their fossil remains are present throughout the geologic record. They are biogeochemical oases that underpin diverse metabolic interactions, transform key nutrients and fix carbon, and can thrive in extreme environments. Mat-forming cyanobacteria can be metabolically versatile and conduct both oxygenic and anoxygenic photosynthesis using sulfide (OP and AP), thereby participating in both oxygen and sulfur cycling. The effect of seasonality on ecological factors constraining photosynthetic production and geochemical cycling in extreme cyanobacterial mats is not well known. In this study, we surveyed the mat community composition via bacterial 16S rRNA genes, microbial activity via metaproteomics, and water physico- and geochemistry over multiple seasons and years of the cyanobacterial mat in Middle Island Sinkhole, an O_2_-poor benthic sinkhole in Lake Huron, Michigan. We found that higher availability of sulfate-rich groundwater, together with higher light intensity, coincided with dominance of the metabolically flexible cyanobacterium *Phormidium* during the summer. Diverse sulfur cycling bacteria were more successful in other seasons when the mat experienced lower light and sulfate availability. These results provide insights into how seasonal environmental dynamics can shape the community structure and metabolisms of microbial mats, ultimately controlling biogeochemical cycling in these ecological hotspots.

## Introduction

The innovation of oxygenic photosynthesis (OP) by cyanobacteria shaped the geosphere and biosphere by driving the rise of oxygen ~2.4 billion years ago (Lyons et al., 2014). In modern systems, cyanobacteria are a core constituent of photosynthetic microbial mats by providing organic matter, fixed nitrogen, and O_2_ for diverse communities (Bolhuis et al., 2014; Stal, 2012). These cyanobacterial products stimulate the metabolism of other microbes such as sulfate-reducing bacteria, sulfide-oxidizing bacteria, and methanogens, which contribute to biogeochemical fluxes of gasses such as methane and sulfide (Hoehler et al., 2001). Some cyanobacteria are also capable of anoxygenic photosynthesis (AP), which uses H_2_S as a reductant in place of H_2_O and produces S^0^ rather than O_2_, and can sustain microbial mats in sulfidic conditions that are toxic to most cyanobacteria (Biddanda et al., 2015; Cohen et al., 1975; de Beer et al., 2017; Dick et al., 2018; Klatt, Meyer, et al., 2016). Sulfidic photic habitats are rare in the modern world but have potential as analogues to understand Earth’s biogeochemical evolution (Dick et al., 2018; Lenton & Daines, 2017; Stal, 2012), and to help address open questions about the biological mechanisms underpinning the pattern of Earth’s oxygenation (Dick et al., 2018; Hamilton et al., 2016).

Light availability exerts strong control on the composition, physiology, and metabolism of cyanobacterial communities. Cyanobacterial niches are defined in part by adaptations to intensity and wavelength of light (Moore et al., 1998; Oberhaus et al., 2007; Ward et al., 2006). Light availability in microbial mats varies on vertical microscales (Jørgensen et al., 1987) that can influence community composition to an extent comparable to variation across larger geographic spatial scales (Al-Najjar et al., 2014). Ice-covered Antarctic lakes illustrate how variation of light due to ice thickness and water depth shapes which cyanobacteria taxa are present in cyanobacterial mats and their productivity, metabolism, and physical structure (Dillon et al., 2020b; Hawes & Schwarz, 2001; Mackey et al., 2017; Moorhead et al., 1997). Cyanobacteria also respond to changes in light at the cellular level via phototaxis (Richardson & Castenholz, 1987) and by modulating their pigment content (Bryant, 1982; Falkowski & LaRoche, 1991) and proteome (Cobley et al., 2002), especially the amount of light-harvesting phycobiliproteins (Hihara et al., 2001).

Light level is also tightly intertwined with the biogeochemistry of cyanobacterial mats because it governs both rates of photosynthesis and the balance of AP and OP, and thereby O_2_ production and H_2_S consumption (Dick et al., 2018; Jørgensen et al., 1986; Klatt et al., 2021; 2020). For cyanobacteria capable of both AP and OP, sulfide is often used as the electron donor before water, thus sulfide-driven AP controls OP such that in the presence of sulfide no OP will take place (Klatt et al., 2015). Higher light levels stimulate higher rates of AP and faster depletion of sulfide; if this outpaces the rate of sulfide supply (*e.g*., from diffusion or local sulfate reduction), then sulfide is exhausted and OP takes over. Hence, oxygenic cyanobacteria may produce oxygen oases in anoxic conditions (Sumner et al., 2015). To date, studies of the effects of light on cyanobacterial mats have focused on geochemical cycling of in situ OP/AP populations on diel scales (Hamilton et al., 2018; Klatt, Meyer, et al., 2016). Nuances observed in diel geochemical cycles can become integrated over longer time scales from seasons to years, and these longer-term patterns are more likely to be recorded in geochemical sediment records. However, the responses of cyanobacterial AP and OP dynamics to seasonal changes in light have not been previously documented.

In addition to light, physical and chemical conditions influence the abundance and activities of microbes in mat systems (Dillon et al., 2020a; Stal, 2012). Sulfide concentration and temperature are key factors in determining the relative abundance of primary producers, which can include cyanobacteria, diatoms, and anoxygenic phototrophs (Camacho et al., 2005; Cohen et al., 1975). In turn, the geochemistry of mats is shaped by the metabolic activities of the consortia of tightly-interacting organisms. Oxygen and sulfide are required and consumed by sulfur-oxidizing bacteria (SOB), and their concentrations influence competition between SOB and cyanobacteria (Klatt, de Beer, et al., 2016). Sulfate-reducing bacteria are often present and active in cyanobacterial mats (Canfield & Marais, 1991; Teske et al., 1998) and may affect cyanobacterial photosynthesis through local production of sulfide (de Beer et al., 2017; Hamilton et al., 2018). Weather-driven events as well as seasonal cycles can shift the delivery of nutrients and light to microbial populations (Bolhuis et al., 2014). Community composition of microbial mats also shifts with season (Cardoso et al., 2019), in some cases due to changes in nutrient availability (Pinckney et al., 1995). In other cases, cyanobacterial mat communities show remarkable resilience to seasonal and climatic change through physiological plasticity (Aguilera et al., 2020; Lionard et al., 2012). The effects of seasonally changing light on the chemistry, composition, and function of redox-stratified microbial mat communities, where light is expected to shape the balance of oxygenic and anoxygenic photosynthesis and thus concentrations of oxygen and sulfide, are largely unknown.

The Middle Island Sinkhole (MIS) in Lake Huron hosts extensive cyanobacterial mat communities that sit at an O_2_/H_2_S interface and switch between oxygenic and anoxygenic photosynthesis depending on supply of sulfide (Kinsman-Costello et al., 2017; Voorhies et al., 2012). The cyanobacterium *Phormidium* dominates the metatranscriptome and metagenome with genes for AP in a variably sulfidic environment and OP in microaerobic conditions (Grim et al., 2021). The mats are bathed in low-oxygen, high-sulfate groundwater that has relatively stable temperature throughout the year (Ruberg et al., 2008), providing an opportunity to study the relationship between seasonal changes in light and groundwater chemistry and microbial community composition and function. We measured light and groundwater chemistry, analyzed microbial community composition via 16S rRNA gene sequencing, and evaluated microbial metabolic activity with shotgun metagenomics and proteomics in samples collected between June and September from 2009 to 2015. We observed strong shifts in the community structure and function of cyanobacteria and sulfur-cycling bacteria, revealing large seasonal impacts on both phototrophs and their metabolic partners.

## Experimental Procedures

### Sample collection

We sampled the microbial community of Middle Island Sinkhole (located at 45° 11.914 N, 83° 19.671 W), a sinkhole of approximately 23 m depth below water surface, 125 m length, and 100 m width in Lake Huron (Baskaran et al., 2016; Kinsman-Costello et al., 2017; Ruberg et al., 2008). We collected samples once in 2009, and one to three times a year between 2011-2015. Scuba divers from the NOAA Thunder Bay National Marine Sanctuary used 20 × 7 cm clear polycarbonate tubes and rubber stoppers to collect intact flat purple mat (**Figure S1**) and sediment cores from the central region of the sinkhole, hereafter referred to as the “arena” (**Figure S2**). Cores were kept upright, in the dark and on ice or at 4°C until they were sampled within 48 hr of collection. Microbial mat was removed intact from cores, homogenized, stored in 2mL microcentrifuge tubes, and frozen at −80°C until DNA or protein extraction.

### Physicochemical measurements

To measure photosynthetically available radiation (PAR) at 23 m depth in the sinkhole, we used a LiCor LI-192 underwater quantum sensor (LiCor Biotechnology) (sampled in 2009-2013, and 2017), a compact optical profiling system for UV light in natural waters (UV C-OPS; Biospherical Instruments Inc.) (Cory et al., 2016) (in 2014), a hyperspectral profiler (HyperPro II profiler, Sea-Bird) (2015-2016), and HOBO loggers mounted 0.25 – 0.75 m above the sediment surface (Onset Computer Corporation) (in 2014-2017). Because HOBO loggers measured light in lux or lumens, their light measurements were calibrated to PAR quantitation through hyperspectral profiling of the 23 m-deep light field at the same time as the loggers. For each month, an average and quartiles of maximum daily light intensity were calculated from the summarized daily maxima and the episodic measurements. *k*-attenuation coefficients were calculated from the linear relationship

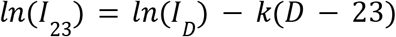

where *I* is irradiance at the last measured depth *D* in meters in vertical profiles obtained through LiCor, C-OPs, and HyperPro II instruments. These *k* values were used to project HOBO-acquired values to uniform depth at 23 m.

Groundwater for conductivity and water stable isotope measurements was collected by divers with 60 mL syringes at specific locations in the sinkhole. Specific conductivity in 2014-2018 was measured using a handheld probe (Yellow Springs Instruments Inc.), as well as calculated from ionic composition. We measured major element anion concentrations in water samples including chloride (Cl^-^), fluoride (Fl^-^), and sulfate (SO_4_^2-^) using electrolytic-suppression ion chromatography (Dionex, Thermo Scientific; average recovery of standards was 10.3%, 10.6%, and 14.1% respectively; standards’ relative percent differences (RPD) were 11.6%, 23.3%, and 4.6% respectively). We measured concentrations of major cation elements including calcium (Ca^2+^), sodium (Na^+^), and magnesium (Mg^2+^) with inductively coupled plasma-optical emission spectroscopy (ICP-OES, Perkin Elmer; average recovery of standards was within 5.2%, 12.6%, and 4.7%, respectively; and standards’ RPDs were 4.8%, 12.7%, and 5.0% respectively). Sulfate measurements are reported from samples collected in 2012-2013 (Kinsman-Costello et al., 2017), as well as samples newly collected in 2014-2018.

We measured stable oxygen (^18^O/^16^O) and hydrogen (^2^H/^1^H) isotope ratios (δ^18^O and δD) in select samples from 2015-2018 to trace the influence of water reservoirs and mixing in the groundwater. We also measured samples from a fountain outside the Alpena, Michigan library in summer and autumn, which putatively taps the same aquifer as the source of MIS groundwater based on its source depth (386 m below surface) (Ruberg et al., 2008). Isotopic measurements were performed on a Picarro L2130-i cavity ringdown spectrometer with an A0211 high-precision vaporizer and attached autosampler (Picarro). The Picarro ChemCorrect software was used to monitor samples for organic contamination. Precision was better than 0.1‰ and 0.3‰ for δ^18^O and δD, respectively, based on replicate injections of deionized water. Data are reported as δ values relative to the Vienna standard mean ocean water normalized to standard light Antarctic precipitation (VSMOW-SLAP) (Coplen 1996). Globally, the relationship between δ^18^O and δD values of meteoric waters is characterized by the Global Meteoric Water Line (GMWL) (Craig 1961), where

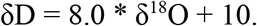

The deviation from global water patterns is labeled the *d-excess*:

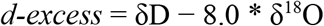

The average *d-excess* for the Great Lakes is +3.2‰, reported as the intercept in the Great Lakes Mean Water Line (GLWL) (Jasechko et al., 2014). δ^18^O and δD values of this study’s water samples were compared to available data from Lake Huron (Jasechko et al., 2014), Michigan groundwaters (Bowen et al., 2012) located within 100km of the sinkhole, and sinkholes offshore of Alpena, including Middle Island Sinkhole (Baskaran et al., 2016). We used the compositions of the Alpena library fountain and surface water samples from Lake Huron as endpoints of a linear mixing model to explain the observed isotopic values in these water samples.

### DNA preparation and sequencing

We extracted up to 0.5 g of wet mat material using a modified version of the MPBio Fast DNA Spin Kit for Soil (MP Biomedical, Santa Anna, CA, USA). In summary, 0.3 g of beads (corresponding to one large bead, seven medium beads, and an equal volume of small beads), sodium phosphate buffer, and MT buffer was used to chemically and mechanically lyse cells, in either the FastPrep instrument for 45 s (samples up to 2013), or horizontal lysis on a vortex mixer for 10 min at speed 7 (2014-2015). After protein precipitation, DNA was cleaned, pelleted, and resuspended in up to 100 uL nuclease-free water. DNA was stored at 4°C for immediate quantification and −20°C for long term.

We used the PicoGreen assay (Invitrogen, Carlsbad, CA, USA) to quantitate double stranded DNA. Samples were diluted to between 1-25 ng/uL and submitted to the University of Michigan Host Microbiome Core for Illumina library preparation and sequencing (Kozich et al., 2013; Seekatz et al., 2015). Bacterial primers 515F/806R were used to amplify the 16S rRNA gene v4 region in a reaction mixture consisting of 5 μL of 4 μM equimolar primer set, 0.15 μL of AccuPrime Taq DNA High Fidelity Polymerase, 2 μL of 10x AccuPrime PCR Buffer II (Thermo Fisher Scientific), 11.85 μL of PCR-grade water, and 1-10 μL of DNA template. Thermocycling was an initial denaturation at 95°C for 2 min, 30 cycles of 95°C for 20 s, 55°C for 15 s, 72°C for 5 min, and a final extension of 72°C for 10 min. PCR products were cleaned and normalized using SequalPrep Normalization Plate Kit (Thermo Fisher Scientific), then quantified and pooled equimolarly according to Kapa Biosystems Library qPCR MasterMix (ROX Low) Quantification kit for Illumina platforms. An Agilent Bioanalyzer kit confirmed library size and purity, and the library pool was sequenced on the Illumina MiSeq using a 500 cycle V2 kit with 15% PhiX for diversity. Sequences can be obtained from NCBI Sequence Read Archive (SRA-SRP067517).

### Quantitative proteomics

Protein abundances in replicate samples from June (n = 3), July (n = 4), and October (n = 3) 2015 were evaluated using diDO-IPTL isobaric peptide labeling (Waldbauer et al., 2017). From 0.25 to 5.0g of wet mat material, proteins were extracted in a denaturing and reducing buffer (1% SDS, 10% glycerol, 10 mM dithiothreitol, 200 mM Tris, pH 8) using heat (95°C, 20 min) and sonication (QSonica Q500). Cysteines were alkylated by addition of iodoacetamide (40 mM, 30 min dark). Following centrifugation to pellet mineral material, proteins were precipitated in acetone (−20°C, overnight) and pelleted by centrifugation. Protein pellets were redissolved in 8M urea and purified by a modified eFASP (enhanced filter-aided sample preparation) protocol (Erde et al., 2014), digested with MS-grade trypsin (Thermo Pierce) at 37 °C overnight, and peptides were eluted and dried by vacuum centrifugation. diDO-IPTL isobaric labeling, wherein peptide N-termini are d_2_/d_0_-dimethylated and C-terminal oxygens exchanged with ^16^O/^18^O-water (Waldbauer et al., 2017), is described in detail at protocols.io (dx.doi.org/10.17504/protocols.io.d2i8cd). An internal standard composed of pooled d_0_/^18^O-labeled aliquots of all samples was used for quantitative comparisons. This standardization normalizes for different bulk sample masses and overall protein extraction/digestion efficiencies. For LC-MS analysis, peptide samples were separated on a monolithic capillary C18 column (GL Sciences Monocap Ultra, 100μm I.D. × 200cm length) using a water-acetonitrile + 0.1% formic acid gradient (2-50% AcN over 180 min) at 360nl/min using a Dionex Ultimate 3000 LC system with nanoelectrospray ionization (Proxeon Nanospray Flex source). Mass spectra were collected on an Orbitrap Elite mass spectrometer (Thermo) operating in a data-dependent acquisition (DDA) mode, with one high-resolution (120,000 *m*/Δ*m*) MS1 parent ion full scan triggering Rapid-mode 15 MS2 CID fragment ion scans of selected precursors. IPTL mass spectral data were analyzed using MorpheusFromAnotherPlace (MFAP; Waldbauer *et al*., 2017). Proteomic mass spectral data are available via ProteomeXchange under accession number PXD031126 and the MassIVE repository (massive.ucsd.edu) under accession number MSV000088706 (username for reviewers, MSV000088706_reviewer; password, a).

We used protein sequences predicted from previously generated metagenomes (Grim et al., 2021; Voorhies et al., 2016) to identify proteins, and metagenome-assembled genomic bins refined through anvi’o (Eren et al., 2015) to link proteins to specific organisms. Briefly, we used ‘anvi-gen-contigs-database’ to profile the metagenomic co-assembly of 15 samples from 2007-2012. The software used Prodigal v2.6.0 (Hyatt et al., 2010) to call genes, and identified single-copy genes belonging to Bacteria (Campbell et al., 2013) and Archaea (Rinke et al., 2013) through HMMER (Eddy, 2011). Commands ‘anvi-import-taxonomy-for-genes’ incorporated gene-level taxonomy calls into the database made using kaiju (Menzel et al., 2016). ‘anvi-profile’ reconciled the mapped reads against the co-assembly to generate differential coverage and tetranucleotide frequency (tnf) information in a profile database, with a minimum scaffold length of 5000 bp. Bins were generated using automated binning with CONCOCT (Alneberg et al., 2014) and MetaBAT (Kang et al., 2019) on contigs not identified as eukaryotic using EukRep (West et al., 2018), and dereplicated using dRep (Olm et al., 2017) and refined using tnf and coverage, yielding 144 metagenome-assembled-genomic bins (MAGs). To recover additional scaffolds that were not long enough for the software-implemented lower limit of 5000 bp, we retained cyanobacterially identified MAGs and used tnf and coverage signals to recover scaffolds that were 2500-5000 bp, 1500-2500 bp, and 1000-1500 bp long. This strategy captured 6011 cyanobacterial contigs that were excluded solely due to length, including contigs with core metabolic genes with potentially >1 copy number such as photosynthetic reaction center gene *psbA*. Our oscillatorial cyanobacterial bins had 70.4% completion/56.3% redundancy (*Phormidium*, 11.0Mbp), 91.6%/42.3% (*Planktothrix*, 5.7Mbp), 95.8%/11.3% (*Pseudanabaena*,4.3Mbp), and 22.5%/1.41% (*Spirulina*, 1.1Mbp), as well as a 527Kbp collection of cyanobacterial scaffolds that were not able to be confidently binned. Additional bins of interest are listed in **Table S1**. Metagenomic scaffolds are available through IMG via genome IDs 3300002026 (5000 bp and longer scaffolds), 3300002027 (1500-4999 bp scaffolds), and 3300002024 (1000-1499 bp scaffolds).

### 16S rRNA gene bioinformatic analysis

Raw pairs of sequencing reads (250 bp) were quality trimmed and merged using ‘iu-merge-pairs’, which is a program in illumina-utils (available from https://github.com/merenlab/illumina-utils) (Eren et al., 2013), using minimum quality score of 25, minimum overlap of 200 bp, and at points of divergence in the overlap the higher quality basecall was retained. Merged reads with five or fewer mismatches were kept for Minimum Entropy Decomposition v. 2.1 (Eren et al., 2014) to generate amplicon sequence variants (ASVs) using the following parameters: -d 4 -N 3 --min-substantive-abundance 5 -V 3 --relocate-outliers. We used GAST (Huse et al., 2008) to call taxonomy using the curated SILVA database, and confirmed with BLASTN against SILVA 123 (Pruesse et al., 2007) (bacteria and archaea), and PhytoRef (Decelle et al., 2015) (chloroplasts). mothur v. 1.33 (Schloss et al., 2009) was used to check for chimeras de novo, and putatively-chimeric ASVs that did not have taxonomy assigned via SILVA 123 and GAST were removed. We searched for sulfate-reducing genera in the Deltaproteobacteria using a taxonomic search for “sulf” or “thio”. The read analysis is outlined here: https://hackmd.io/s/r1CGeQs_G

### Statistical analyses

We used the R statistical environment (R Core Team, 2015) in RStudio (RStudio Team, 2014) to analyze ASVs and proteins. For environmental samples, we used Morisita-Horn metric to calculate a distance matrix on Hellinger-transformed bacterial relative abundances, as input for nonmetric multidimensional scaling (NMDS) with autotransformation = TRUE. For correlation network analyses, we retained bacterial genera that were at least 0.1% of the total community (n = 67). We generated one network using rcorr on a Bray-Curtis distance matrix calculated from the read counts of genera, and after a Benjamini-Hochberg false discovery rate correction, we retained all correlations p < 0.001. We used CoNetinR (Faust & Raes, 2016) to generate three additional network matrices using Spearman’s rho correlation, Pearson correlation, and Bray-Curtis, implementing the ReBoot procedure as described in the original CoNet (Faust & Raes, 2012) and Benjamini-Hochberg FDR correction, to keep correlations p < 0.001. The final correlation network contained ASVs and edges whose direction (positive or negative) were supported by all four matrices, with the edges representing the average of the measures. Using such statistical processing likely reduces the impact of taxonomic bias skewing differential abundance analyses (McLaren et al., 2022). The final network was visualized in Gephi and in a correlogram via corrplot, with the standard deviation of the mean in the lower triangle. We used amova (Excoffier et al., 1992) in the R package *pegas* (Paradis, 2010) for testing significant difference in bacterial community structure between months and seasons, and LEFSE in the Galaxy Project (Afgan et al., 2018) to identify taxa as biomarkers for seasons and months; those taxa are indicated with significance values. For evaluation of differential abundance of proteins between seasons, we calculated the weighted mean and weighted standard deviation of the log2-normalized abundance ratios of samples taken from the same month. Paired t-tests (p < 0.05) corrected with Benjamini-Hochberg false discovery rate were used to retain significantly different weighted mean abundances.

## Results

### Seasonal light dynamics

We measured PAR over several years with varying frequency and methods. *k*-extinction coefficients for PAR ranged from 0.12 to 0.14, were not related to season, and were consistent with previous measurements in oligotrophic Lake Huron (Yousef et al., 2017). Although light measurements were taken throughout the year, observations of the microbial mats were limited to months May-June (hereafter referred to as “spring”), July-August (“summer”), and September-October (“autumn”). Within the timeframe of our mat observations, PAR was generally highest in July and lowest in May and October (**Figure 1**). In addition to the expected astronomical variation of solar radiation, these measurements also reflect varying turbidity in the water column due to phytoplankton growth, and episodic shading from clouds and/or ice.

**Figure 1.**
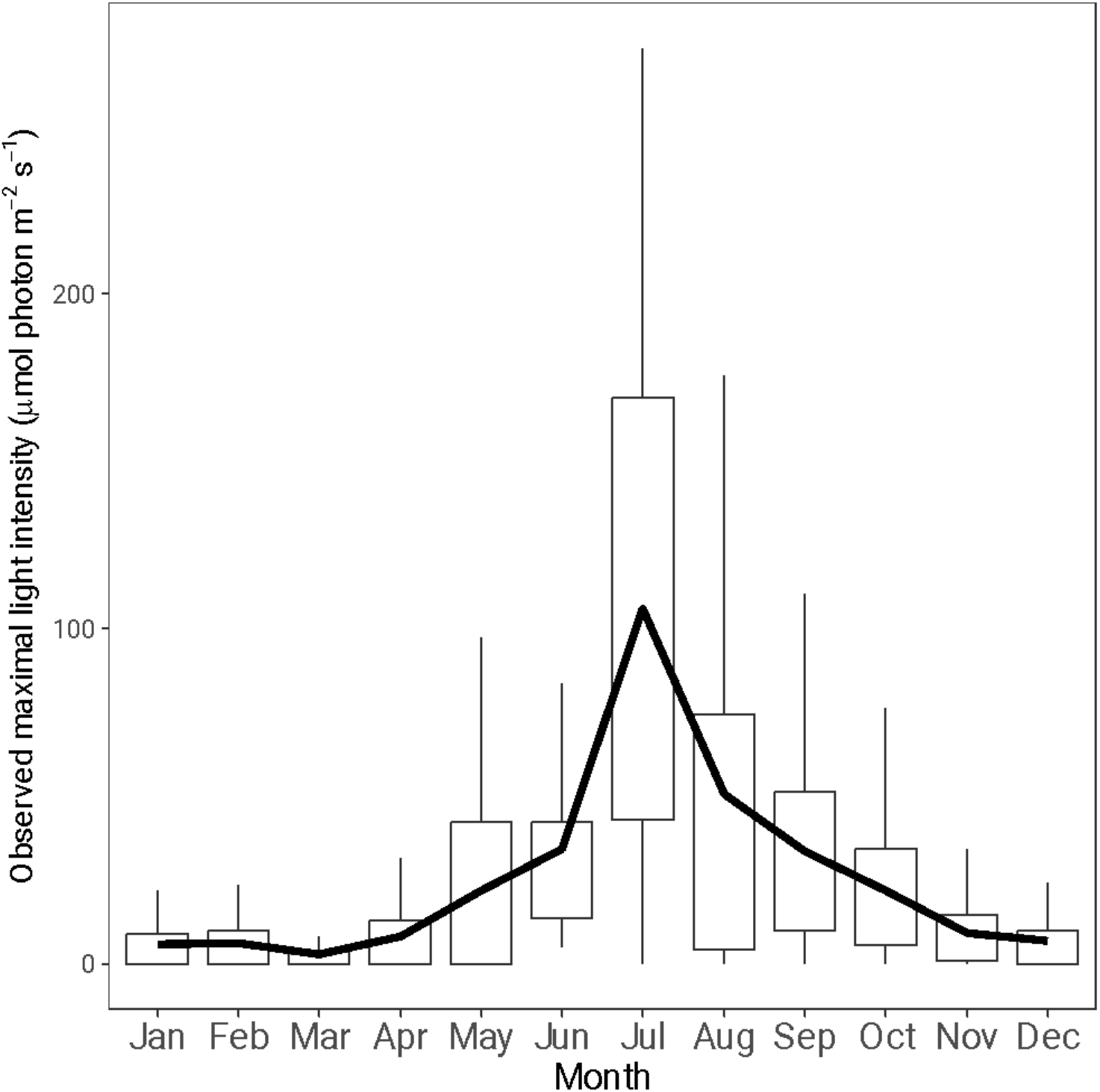
Maximal daily light observed each month in the sinkhole above microbial mats. For each month, daily maximum light values from logged data at 23 m depth (2014-2017) and recorded light from episodic measurements to 23 m depth (2009-2016) (grey transparent circles) were summarized in box and whisker plots and averaged per month. Boxes represent the 25-75^th^ percentiles with whiskers covering the minimum and maximum values. The average is plotted with the thick line.

The quality of light also varied between seasons. Seven hyperspectral profiles of wavelengths 348-801 nm showed that at 23 m depth in the MIS arena, blue-green light was most available (**Figure S3**). As expected, the amount of blue and green light was directly related to overall PAR availability, whereas red light was generally below 0.002 μmol m^-2^ s^-1^ nm^-1^. We observed subtle differences in availability of other wavelengths of light (**Figure S4**); the average attenuation coefficient for green light (530-570 nm) in July (average of four casts: k = 0.109 ± 0.007 m^-1^) was generally higher than in June (average of 2 casts *k* = 0.099 ± 0.009 m^-1^) and October (1 cast, *k* = 0.103 m^-1^) (**Table S2**), indicating green light is more rapidly attenuated in summer than in the other seasons. Hyperspectral profiles (n = 2) at an open water site outside of the sinkhole (Kinsman-Costello et al., 2017) in June 2015 were comparable to those in the sinkhole, indicating similar water column dynamics between the two sites.

### Seasonal variation of groundwater chemistry

Groundwater that vents from its main source, the sinkhole alcove, is thermally and chemically distinct from overlying Lake Huron water, and as it flows through the sinkhole it slowly mixes with lake water (Baskaran et al., 2016; Ruberg et al., 2008). The conductivity of the dominant source of groundwater (hereafter referred to as “alcove groundwater source”) was on average 3055 μS cm^-1^, and it attenuated to an average of 1909 μS cm^-1^ as it distributed in the bowl-shaped arena of the sinkhole (hereafter referred to as “MIS arena water”) (**Table S3, Figure S5**). These values are higher than lake surface water (208 μS cm^-1^), similar to previously reported values for MIS (Baskaran et al., 2016), and lower than the Alpena fountain, which averaged 3624 μS cm^-1^ and is thought to reflect the chemistry of source groundwater without any mixing.

The isotopic composition of water samples from the MIS alcove and arena was depleted in both ^18^O and D compared to lake surface (**Table S4, Figure 2**). Isotopic composition of surface waters sampled in this study were indistinguishable from previous measurements of Lake Huron water samples (**Figure 2A**) (Jasechko et al., 2014). Additionally, Alpena fountain and alcove groundwater samples were isotopically similar to groundwaters within 100km of the sinkhole (Bowen et al., 2012) and submerged sinkholes in the immediate area (Baskaran et al., 2016). Year-round δ^18^O values for surface waters averaged (± 1 standard deviation) −7.1 ± 0.1‰, whereas the Alpena fountain averaged −12.2 ± 0.1‰. More seasonal variability was observed in δ^18^O measurements from the sinkhole arena and alcove. In arena water samples, the average δ^18^O in spring, summer, and autumn were −9.1 ± 0.4‰, −10.0 ± 0.3‰, and −9.6 ± 0.6‰, respectively. In the alcove, across the respective seasons δ^18^O averaged −11.4 ± 0.1‰, −11.5 ± 0.1‰, and −10.7 ± 0.8‰. The same regional and seasonal signals were observed in δD measurements. The Alpena fountain samples had substantially lower δD (average: −84.2 ± 1.7‰) compared to the lake surface samples (average: −52.9 ± 0.7‰). Seasonal averages for δD values in arena water samples were −64.8 ± 2.3‰ (spring), −70.1 ± 1.3‰ (summer), and −67.1 ± 3.4‰ (autumn). Alcove water samples had even lower δD values, with seasonal averages of −77.9 ± 0.4‰ (spring), −78.7 ±0.4‰ (summer), and −74.3 ± 4.7‰ (autumn).

**Figure 2.**
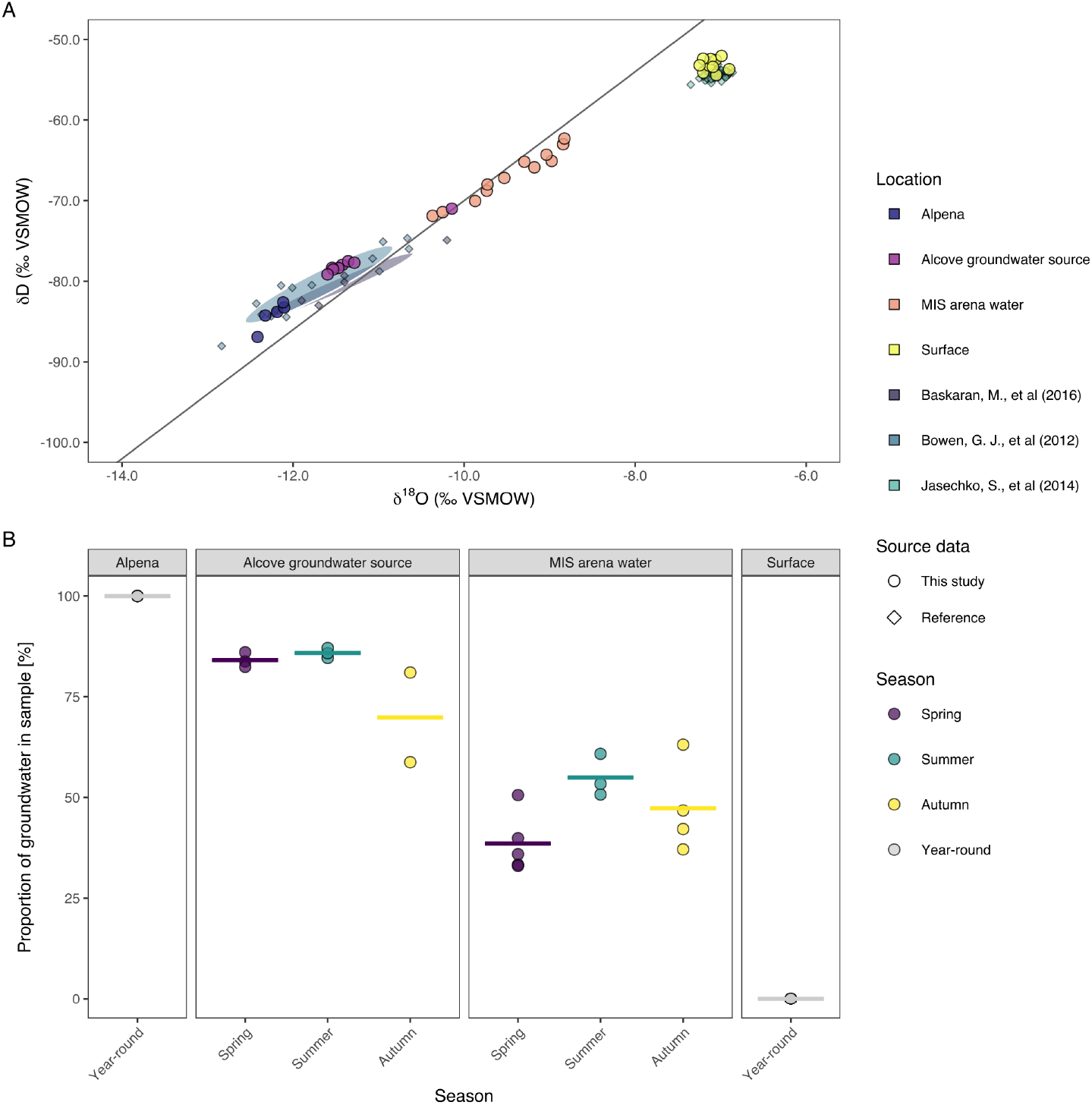
Measured values of δD and δ^18^O and estimated proportion of groundwater in water samples. A. Isotopic compositions of samples from this study are plotted as circles and colored by location. Samples from other references are plotted as diamonds, and colored by study. B. The average δ^18^O values for Alpena fountain and surface were used as end-members in a linear mixing model to estimate the proportion of ground water in each sample. Isotopic compositions of samples from this study are plotted as circles and colored by season. The horizontal line represents the average proportion of groundwater in each season per location.

Isotopic compositions can help constrain water sources and influences. In this study, the average *d-excess* of surface water samples was +4.2 ± 0.6‰, which is slightly higher than the previously reported Lake Huron-specific average *d-excess* of +2.6 ± 1.0‰ but within the range of measurements for the Great Lakes (**Figure S6**) (Jasechko et al., 2014). Given the sinkhole’s proximity to shore, the slightly elevated *d-excess* values in surface waters likely reflects incomplete mixing of riverine input (*d-excess* values of rivers in Lake Huron catchment: +11.3 and +12.5‰) and terrestrial runoff (Jasechko et al., 2014). In contrast, *d-excess* was highest for Alpena fountain samples (average 13.7 ± 0.8‰), which aligns with the *d-excess* values of terrestrial groundwaters in Michigan’s Lower Peninsula most impacted by lake-effect precipitation (Bowen et al., 2012). High *d-excess* values in the aquifer that sources Alpena fountain (386 m depth below surface) suggest a large contribution of lake-effect precipitation.

We leveraged the δ^18^O and δD data to constrain water sources to Middle Island Sinkhole by constructing a linear mixing model with two isotopic endmembers: the lake surface water samples (no groundwater), and the Alpena library fountain (representing the groundwater reservoir). In the sinkhole arena, the model estimates the proportion of water sourced from the groundwater aquifer increases from spring to summer (p < 0.03 by Kruskal-Wallis testing) and decreases summer to autumn (p < 0.06) (**Figure 2B**). We observed a similar (though not significant) trend in the alcove source waters, and corroborated the linear mixing model using ion concentrations (SO_4_^2-^, Cl^-^, Na^+^) (**Figure S7**). Water column mixing in spring and autumn can lead to a larger input of lake surface water to the sinkhole, yielding higher values of δ^18^O and smaller *d-excess* in the arena. In the summer, lower δ^18^O and larger *d-excess* values in sinkhole arena and alcove waters point to a greater contribution of the groundwater aquifer, likely due to thermal stratification in the water column isolating the cold groundwater layer from surface water.

While the seasonal signal is strong and robust, sampling location may also influence isotopic variability in MIS water. Samples in 2016 were collected in georeferenced locations (**Figure S2**), whereas in 2015 they were collected opportunistically with sinkhole mat sampling, and in 2017-2018 at one consistent location. Focusing on the georeferenced locations from 2016, the mixing model of isotopic composition suggests that the proportion of aquifer groundwater decreases with greater distance from the alcove source (**Figure S2**). There are small local sources of groundwater around the circumference of the sinkhole arena (Baskaran et al., 2016), yet the largest source and dominant influence on the isotopic composition is from the alcove groundwater source. Turbulence between the groundwater layer and overlying lake water, and increased distance from the alcove source likely attenuate the signal of aquifer groundwater in arena water samples that are farther from the alcove, yet they are still chemically distinct from surface water and other depths sampled in Lake Huron (Jasechko et al., 2014).

### Microbial community structure varies seasonally

We evaluated the bacterial community observed in the microbial mat samples from 2009-2015. Samples were categorized as “spring” (from May and June, n = 9), “summer” (July and August, n = 13), and “autumn” (September-October, n = 12). Diatom chloroplast 16S rRNA genes ranged in abundance from 0.14-44% (average: 12.4% ± 12.7) of all sequencing reads, belonged primarily to Bacillariophyta;Cymbellaceae, and were removed from further analysis due to the highly variable abundance of 16S rRNA gene copies in their cells (Green, 2011).

Bacterial communities were dissimilar by month (AMOVA, p < 0.05) and by season (p < 0.08), with spring clustering distinctly from autumn, and summer communities showing a degree of mixing between the two (**Figure 3, Figure S8, Table S5**). Seasonal trends were also apparent in specific taxa, including the dominant cyanobacteria, *Phormidium* and *Planktothrix* (**Figure 4**). *Phormidium* was more abundant in summer samples (on average 33 and 35% of the total bacterial community in spring and summer respectively) compared to autumn samples (4.2%) (LEFSe, p < 0.05), whereas *Planktothrix* is more frequently observed in autumn and spring samples (8.4-8.0% compared to 1.9% in summer) (LEFSe, p < 0.05) (**Table S5**). Other cyanobacterial taxa were typically 5% or less of the community.

**Figure 3.**
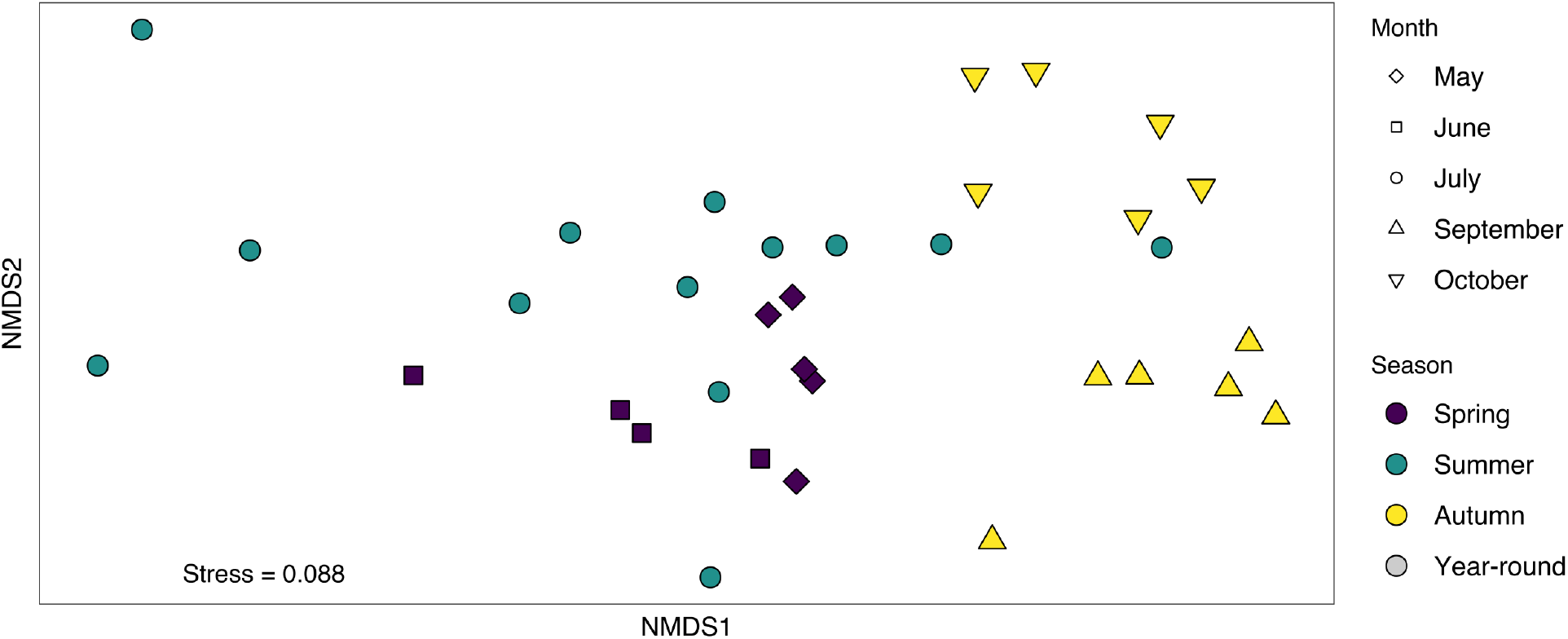
Nonmetric multidimensional scaling plot of bacterial community structure by season, using Morisita-Horn distance matrix calculated from Hellinger-transformed bacterial relative abundances. Samples in May (diamonds) and June (squares) are colored purple, July samples are green circles, and samples from September (upright triangles), and October (lower triangles) are colored yellow.

**Figure 4.**
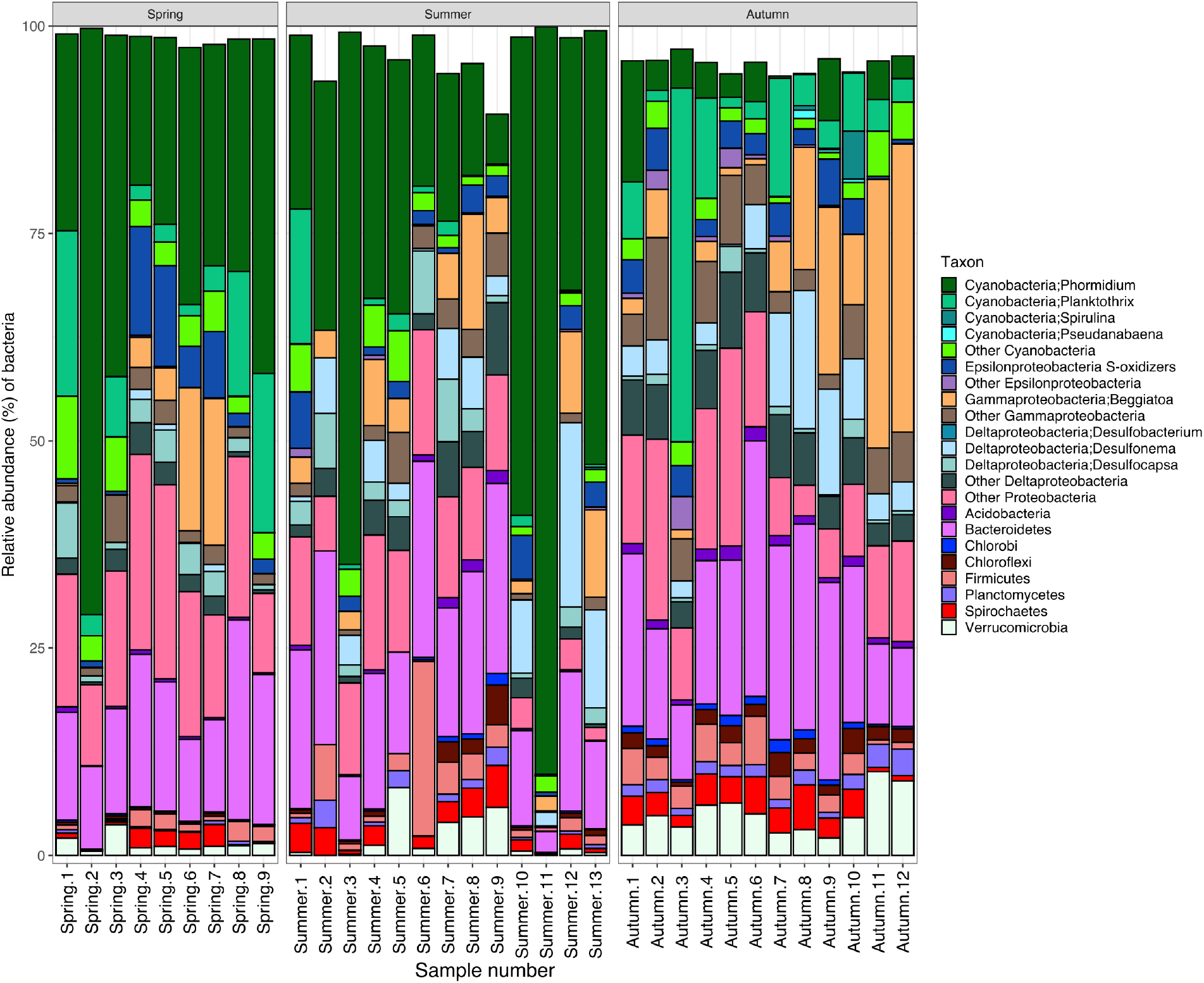
Relative abundance of relevant bacterial taxonomic groups in samples, grouped by season. Sample naming on x-axis reflects chronological order within seasons. Relative abundances of key genera were summed and presented, and classes and phyla without those genera are represented.

Deltaproteobacteria, including putative sulfur and sulfate reducing bacteria (hereafter “SRB”) such as *Desulfonema* and *Desulfocapsa*, ranged from 4.8% on average of the total bacterial community in spring samples to 13% in summer and autumn (**Table S6**). Within this group, however, we observed some seasonal peaks in specific genera. Many taxa followed the pattern of *Desulfonema*, which was more abundant in summer and autumn (5.9-6.0% compared to 0.34% in spring, LEFSe, p < 0.05). In contrast, *Desulfocapsa* is more abundant in summer samples compared to autumn (average 2.7-3.0% versus 0.93%). Another SRB, *Desulfotalea*, is more abundant in spring than later months (0.0012% vs 0.00061% in summer and <0.0001% in autumn).

The most abundant putative sulfur-oxidizing bacteria (hereafter “SOB”), identified by classification in taxonomic groups known to be SOB, were primarily gammaproteobacteria (*Beggiatoa)* and 5 groups of epsilonprotebacteria (*Arcobacter*, *Sulfurospirillum*, *Sulfurovum*, *Sulfuricurvum*, and *Sulfurimonas*). Across seasons, *Beggiatoa* was at least as abundant (4.8-11%) as putative SOB epsilonproteobacteria combined (2.4-4.8%). Other gammaproteobacteria and epsilonproteobacteria comprised 0-5% of the community on average. Other proteobacterial members and Bacteroidetes each constitute 12-18% on average throughout the year. Firmicutes, Spirochaetes, and Verrucomicrobia were each on average 5% or less of the bacterial community. Other bacterial groups, including Acidobacteria, Chloroflexi, Chlorobi, and Planctomycetes, were infrequently observed (<1% typically).

### Network analysis identifies taxa with correlated abundances

We used correlation network analysis to understand potential positive and negative relationships between the abundances of 60 bacterial genera in the microbial mat (**Figure 5, Figure S9**). All relationships evaluated were positive in direction and had r ≥ 0.50. *Beggiatoa* was solely correlated with *Desulfonema*, which was also correlated with *Paludibacter* (Bacteroidetes) and *Desulfobacterium*. *Sulfuricurvum* was linked to other putative sulfide-oxidizing taxa (*Sulfurospirillum*, *Sulfurovum*) as well as the putative SRB *Desulfobacterium*. *Sulfurovum* in turn linked *Sulfurimonas*, *Desulfocapsa*, *Desulfobulbus*, *Desulfomicrobium*, and *Desulfobacula* to the group. *Thiothrix*, another sulfide-oxidizing gammaproteobacterium, was correlated with *Desulfomicrobium*. *Planktothrix*’s sole correlation was with *Paludibacter*. *Phormidium* was not correlated with any taxa.

**Figure 5.**
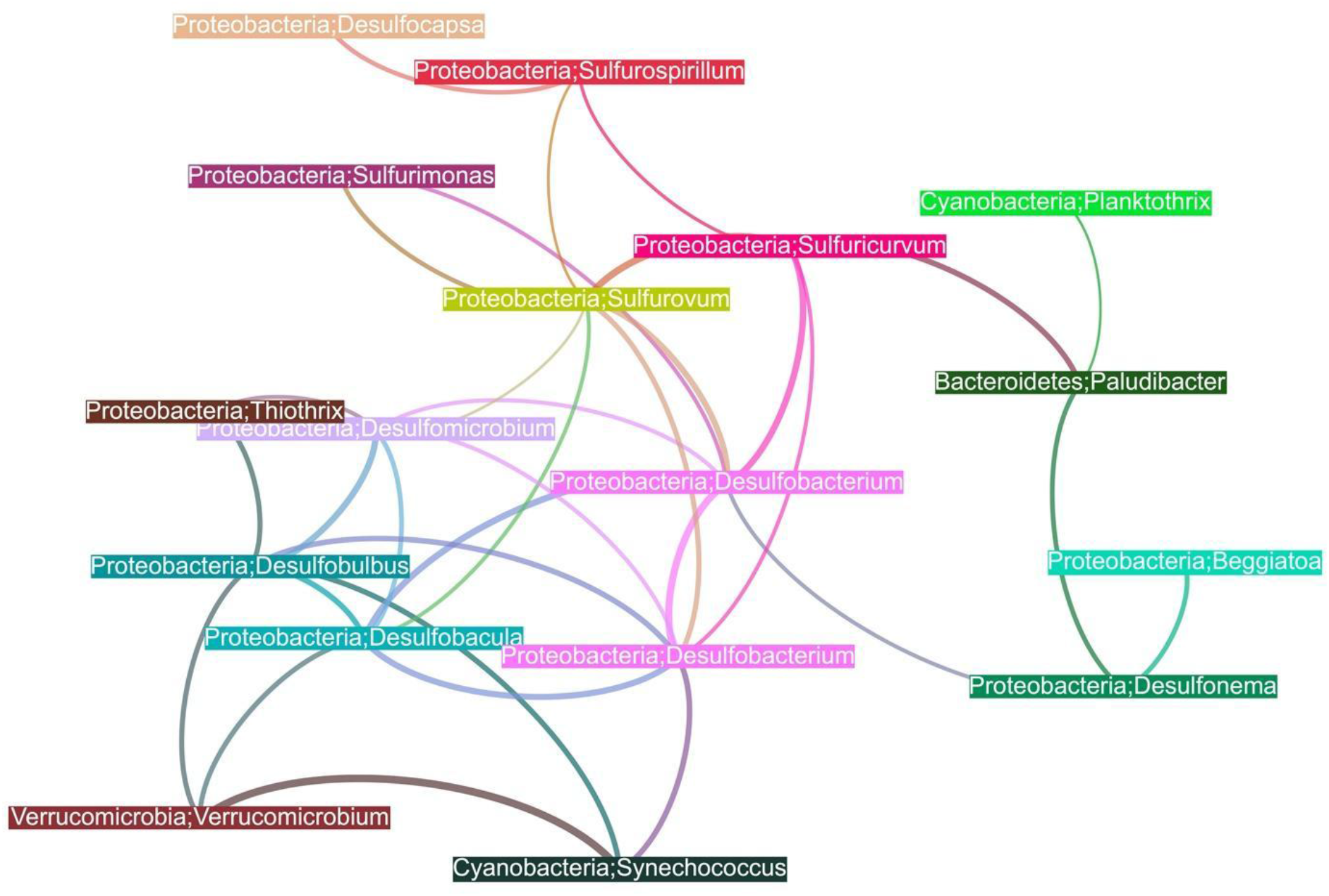
Correlation network between relative abundances of key genera. Bray-Curtis distance matrix was calculated from read counts of genera and used to generate a correlation network. Only significant (p < 0.001) positive (r ≥ 0.6) correlations between specific organisms are plotted, and line thickness conveys strength of the correlation. Colors are arbitrary, and line colors are blends of the nodes they connect.

### Community function differs by season

Quantitative proteomics analyses showed that the abundances of specific proteins belonging to specific genomic bins varied across the seasons (**Figure 6**, **Table S7**). Nearly half (317/789) of proteins identified belonged to cyanobacteria, of which *Phormidium* was the dominant contributor (215 proteins). Across the entire proteomic dataset, 54% of observed spectra were attributed to phycobilisome-related proteins, with most from *Phormidium* (62%). Based on genome size, redundancy, and completion metrics, there were likely multiple *Phormidium* strains in our MAG bin (**Table S1**). We were also unable to resolve several observed proteins to specific MAG bins (referred to as “unknown organism”). When possible, proteins were assigned taxonomy based on taxonomic profiles of genes from the source contig.

**Figure 6.**
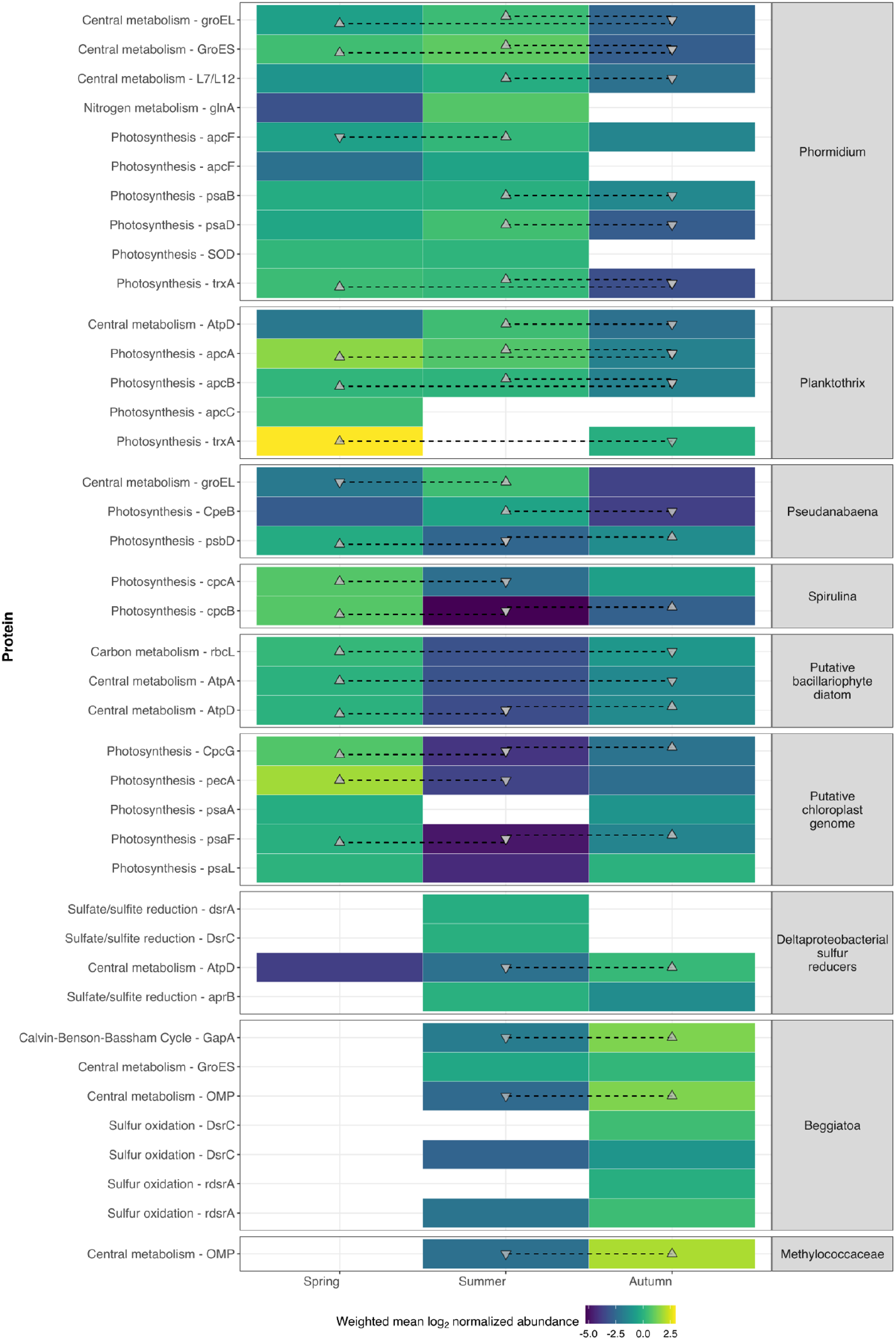
Significantly differentially abundant proteins observed in each season. Weighted-mean log_2_-normalized observations of proteins were used in paired t-tests to determine significant changes in abundance. Significant differences in abundances between two season units are indicated with a line between them and the direction of the triangles. For example, a protein significantly abundant in spring versus summer is indicated with an upright triangle in the spring column linked to an upside-down triangle in the summer column. Proteins are grouped by MAG bin, and described by function.

*Phormidium* dominated the proteins assigned to genome bins in spring and summer (1076/1960 spectra in June and 4269/5535 spectra in July), whereas diatom proteins (Bacillariophyta) were most abundant in autumn (376/1796 spectra observed in September). Despite the dominance of *Phormidium* photosynthetic proteins in the dataset, only a handful varied significantly in abundance between seasons (**Table S7**). We observed photosystem I proteins PsaB and PsaD in significantly lower abundance in autumn compared to summer. Thioredoxin protein TrxA was significantly less abundant in autumn. An allophycocyanin subunit ApcF protein was less abundant in spring compared to summer. Other phycobilisome proteins composing allophycocyanin, phycoerythrin, and phycocyanin had dynamic abundances across the seasons (**Table S7**).

*Phormidium* proteins related to growth also varied in abundance between seasons. Chaperonins GroEL and GroES were significantly less abundant in autumn compared to the other seasons. A ribosomal protein L12 RplL was significantly lower in autumn than in summer. Superoxide dismutase (SOD) and a rhodanese involved in sulfur cycling were abundant exclusively in summer and not observed in other months. Although these SOD and rhodanese proteins are on the same contig as a microaerobic PsbA and a type B sulfide-quinone reductase (Grim & Dick, 2016), we were not able to detect SQR, the sulfide quinone reductase responsible for anoxygenic photosynthesis in cyanobacteria. Further, peptides for *Phormidium*’s photosystem II protein PsbA were not recovered from these samples.

Additional phototrophs, including *Planktothrix*, *Pseudanabaena*, *Spirulina*, other unidentified cyanobacteria, and diatoms, also had photosynthesis- and growth-related proteins that were differentially abundant between seasons, generally higher in summer compared to autumn (**Figure 6, Table S7**). This included subunits of accessory pigments allophycocyanin, phycocyanin, phycoerythrin, phycocyanin, and phycoerythrocyanin, thioredoxin, and F0F1-type ATP synthase. Only a few proteins from phototrophs were more abundant in autumn, including a diatom’s alpha subunit of F0F1 ATP synthase. Spectral abundance of proteins from *Phormidium* peaked in summer (58.4% of those months’ recovered spectra) while *Planktothrix* was responsible for more observed spectra in spring (13.4% of those months’ spectra) than in summer and autumn (6.4% each). We observed more MS/MS spectra belonging to diatom PsbA (23 across the dataset) compared to PsaA (6).

Proteins from sulfur-oxidizing bacteria were generally more abundant in autumn and in some cases only detected in autumn (**Table S7**). This included *Beggiatoa* proteins for sulfur oxidation (dissimilatory sulfite reductase subunits DsrC and RdsrA), glyceraldehyde 3-phosphate dehydrogenase GapA, and an outer membrane protein. Sulfate-reducing versions of dissimilatory sulfide reductase (DsrA and DsrC) were observed only in summer, and a beta subunit of adenosine-5’-phosphosulfate reductase for dissimilatory sulfate reduction from Desulfobacteraceae was less abundant in autumn compared to summer (**Table S7**). In contrast, a beta subunit of F0F1 ATP synthase from *Desulfotalea* was significantly more abundant in autumn compared to summer months.

## Discussion

Cyanobacterial oxygenic photosynthesis produces O_2_ and fixed carbon, providing an energetic and biogeochemical foundation for networks of interacting organisms in microbial mats. Cyanobacterial mats exposed to sulfide also perform anoxygenic photosynthesis, producing oxidized sulfur compounds, with the balance of oxygenic and anoxygenic photosynthesis being sensitive to irradiance intensity and having major impacts on mat biogeochemistry (Klatt et al., 2015). Here we studied how seasonal changes in light and water chemistry affect microbial community composition and metabolic functions in cyanobacterial mats from Middle Island Sinkhole (MIS) over several years. We found that the composition of the microbial community and the proteome of key microbes such as the dominant cyanobacterium (*Phormidium*), the dominant sulfide-oxidizing bacterium (*Beggiatoa*), and a variety of sulfate/sulfur reducing bacteria shifted between seasons (**Figure 7**). These results are consistent with the hypothesis that seasonal changes in light availability impact sulfur cycling, with more intense sulfur cycling in autumn, when lower light levels should promote more anoxygenic and less oxygenic photosynthesis (Klatt et al., 2015; 2021).

**Figure 7.**
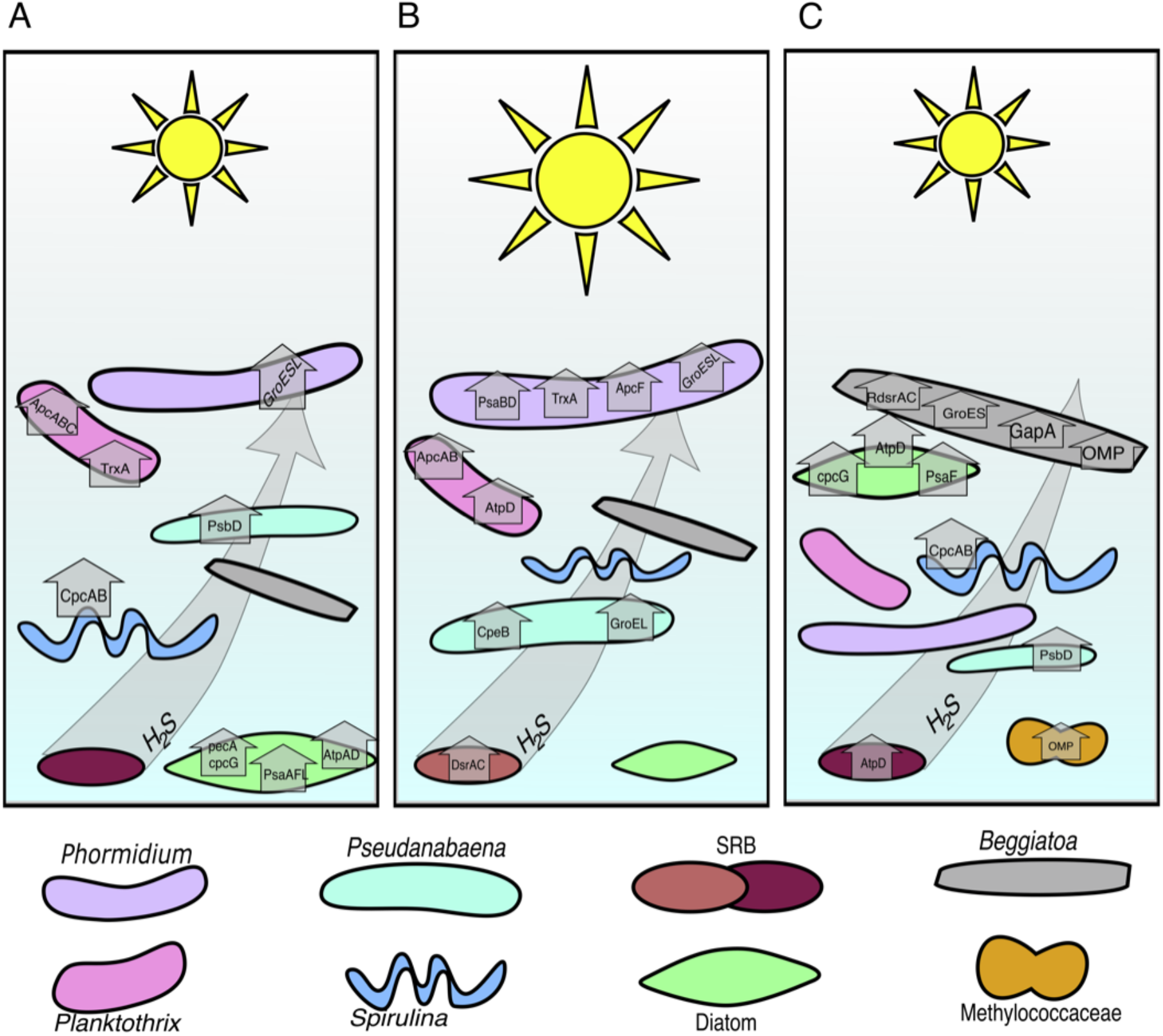
Seasonal changes in the functions and membership of the cyanobacterial mat in MIS. In spring (A), the microbial community is dominated by cyanobacteria able to thrive at low light (*Planktothrix*, *Pseudanabaena*, and *Spirulina*). In summer (B), though *Planktothrix* and *Pseudanabaena* are still active, *Phormidium* is the most abundant and active cyanobacterium. Different sulfate-reducing bacteria (SRB) are also active in summer and autumn (C). *Beggiatoa* and Methylococcaceae are abundant and active in autumn. Diatoms are present throughout the year, but their proteins are more abundant in low-light autumn and spring. Vertical placement of microbial members reflects relative depth in the microbial mat.

We characterized environmental factors that may drive these shifts, focusing on light and groundwater chemistry which directly influence photosynthesis and sulfate delivery. By altering rates of oxygenic and anoxygenic photosynthesis and sulfate reduction, and the abundance of taxa conducting these processes, light and groundwater chemistry are expected to indirectly affect available organic carbon and electron donors (H_2_S) and acceptors (O_2_ and sulfate). The temperature of groundwater bathing the mats remains 8-10°C through the year, with only transient changes from mixing during storm events (Ruberg et al., 2008). Photosynthetically active irradiance (PAR) and spectral quality of irradiance varied seasonally (**Figures 1, S3, S4**); with green light (which phycoerythrin maximally absorbs) most available in spring and least available in summer likely due to growth of planktonic cyanobacteria in the overlying water (Fahnenstiel & Carrick, 1991). These trends in light quality and quantity are expected to promote modulation of phycoerythrin (which maximally absorbs green light) but not phycocyanin (which maximally absorbs red light) by cyanobacteria, known as group II type complementary chromatic acclimation (De Marsac & Houmard, 1993). These dynamics in light intensity and quality are also likely to differentially affect ecotypes adapted to different light niches (Stomp et al., 2004).

The isotopic and specific conductivity measurements show that the benthic water layer in the sinkhole is composed of seasonally-changing proportions of Lake Huron surface water and venting sulfate-rich groundwater. The groundwater source is chemically distinct from the lake surface, and isotopic composition is invariant over the years. Our linear mixing model suggests that groundwater supplies a substantial amount of sulfate to the sinkhole. Over the course of the year, groundwater discharge is higher in summer compared to spring and autumn. Sulfate measurements in overlying groundwater and previously collected samples (Kinsman-Costello et al., 2017) fit reasonably with predictions from the linear mixing model.

The higher abundance of *Phormidium* in spring and summer than in autumn (**Figures 4, 6**) followed light levels and was also reflected in the lower abundance of *Phormidium* proteins for allophycocyanin and photosystem I proteins PsaB and PsaD in autumn. Superoxide dismutase was present in spring and summer but not in autumn, suggesting oxidative stress associated with higher light levels. Seasonal changes in the abundance of other phycobiliproteins were more complex. The *Phormidium* genome bin contained multiple copies of genes for phycoerythrin and phycocyanin, which could result from multiple copies in the genome and/or heterogeneity between strains (Guan et al., 2007; Rinalducci et al., 2008) within the bin (**Table S1**). The abundance of the different isoforms of these proteins varied through the season (**Figure 6),**consistent with complementary chromatic acclimation in response to changing light conditions (De Marsac & Houmard, 1993). Variation of phycocyanin and phycoerythrin abundance has also been implicated in photophysiological acclimation to different light levels in nearby shallow water cyanobacterial mats with similar microbial communities (Snider et al., 2017).

Seasonal differences in the *Phormidium* proteome also indicated broader changes in its physiology. The higher abundance of *Phormidium* ribosomal proteins and GroES/EL proteins in summer likely reflects higher growth rates (Georgopoulos & Welch, 1993) (Jahn et al., 2018). Reduced metabolic activity in autumn was also indicated by lower thioredoxin levels. Light energy from photosystem I is used to reduce thioredoxin (TrxA), which interacts with enzymes in the Calvin cycle and oxidative pentose phosphate cycle to balance carbon fixation and catabolism (De Marsac & Houmard, 1993). By moderating levels of reduced TrxA, cyanobacteria control key cellular and metabolic processes, such as carbon fixation, nitrogen metabolism, oxidative stress response, and light harvesting (Blankenship, 2014), and expression of *trxA* transcripts is reduced at lower light levels (Navarro et al., 2000; Pérez-Pérez et al., 2009).

In contrast to the reduced abundance of *Phormidium* in autumn, *Planktothrix* was more abundant in spring and autumn than in summer, suggesting a degree of niche partitioning between these dominant cyanobacteria. The higher abundance of *Planktothrix* in lower-light seasons is consistent with its tolerance to low light levels (Walsby & Schanz, 2002). Further, *Planktothrix* proteins for phycobilisomes and thioredoxin were significantly higher in spring, suggesting higher photosynthetic activity at that time.

In addition to light, sulfide has a strong impact on the physiology of cyanobacteria (Dick et al., 2018), and influences community composition. Seasonal influence of sulfide on MIS mat communities is likely given evidence for stronger sulfur cycling in autumn, including the increased abundance and activity of sulfur cycling bacteria presented here as well as previous measurements of higher sulfide levels in autumn (Kinsman-Costello et al., 2017). *Phormidium* and *Planktothrix* both have genes for sulfide quinone reductase (*sqr*), which oxidize sulfide for the purpose of anoxygenic photosynthesis and/or detoxification (Dick et al., 2018; Voorhies et al., 2012). Our proteomic dataset contained neither SQR proteins nor those from variant photosystem II (*psbA*) genes optimized for low oxygen conditions (Dick et al., 2018; Grim & Dick, 2016), which may have been masked by the much more abundant phycobilisome proteins (which can constitute 40% of cell protein) (Grossman et al., 1995). Lack of protein detection does not imply the absence of function; for example, *Phormidium’*s PsbA was not detected yet we have verified oxygen evolution in the mat (Klatt et al., 2021). However, missing the SQR proteins does preclude inference of the balance of OP and AP photosynthetic modes of these dominant cyanobacteria in the MIS mat, and their relationship to the observed changes in community composition over the seasons.

The increased abundance of sulfur-oxidizing bacteria and sulfur/sulfate-reducing bacteria in autumn suggests an intensification of the sulfur cycle, which has implications for primary production and O_2_ cycling in the cyanobacterial mats (**Figure 7**). *Beggiatoa* is a dominant sulfur oxidizer in MIS mats, contributing up to 35% of bacterial sequences. It was especially abundant in autumn, when we observed higher abundances of key *Beggiatoa* proteins, such as glyceraldehyde 3-phosphate dehydrogenase involved in the Calvin cycle, an outer membrane protein likely involved in motility (Yu & Kaiser, 2006), and reverse dissimilatory sulfite reductase proteins involved in sulfur oxidation. Lower light levels, as observed in autumn at MIS, also suppress anoxygenic photosynthetic sulfide oxidation, making sulfide more available for chemolithotrophs (Klatt, Meyer, et al., 2016). Taken together, lower light and higher sulfide levels in autumn (Kinsman-Costello et al., 2017), along with increased proteomic and cellular abundance of *Beggiatoa*, suggests that seasonal changes in light availability exerts strong influence on the abundance of sulfide and sulfur-oxidizing bacteria. *Beggiatoa* is motile and moves vertically to position itself within the gradients of sulfide and oxygen at MIS (Biddanda & Weinke, 2020) and other redox-stratified microbial mats (Jørgensen & Revsbech, 1983). This migration also shades the cyanobacteria on the mat surface, reducing photosynthesis and O_2_ production (Dick et al., 2018; Klatt et al., 2021; Klatt, Meyer, et al., 2016). Thus, the seasonal shift towards lower light and more sulfur-oxidizing bacteria likely hastens declines in rates of photosynthesis and O_2_ production.

The 16S rRNA gene and proteomic data indicate that SRB abundance also increased in autumn and suggest interactions with *Beggiatoa* via the sulfur cycle. Network analysis of the 16S rRNA gene time series showed correlated relative abundances of *Beggiatoa* and *Desulfonema*,the most abundant SRB we observed, likely reflecting a metabolic interaction in which sulfide produced through sulfate reduction by *Desulfonema* is used by *Beggiatoa* as an electron donor, as observed in other cyanobacterial mats (Klatt et al., 2020; Stal, 2012). The higher overall abundance of deltaproteobacterial SRB in summer and autumn (**Table S6**) is consistent with higher sulfide availability in the mat and sediment later in the year (Kinsman-Costello et al., 2017). However, other putative SRB taxa, such as *Desulfocapsa* and *Desulfotalea*, showed a distinct seasonal pattern, with higher abundance in spring. Proteins from unbinned sequences involved in sulfate reduction (DsrA, DsrC, ApsR) were observed in summer samples either exclusively or in greater abundance. Thus, SRB taxa are differentially abundant across the seasons, and they are also differentially abundant with depth in mats and underlying sediments (Kinsman-Costello et al., 2017), suggesting that SRB partition niches across both time and space. In addition to sulfate reduction, *Desulfocapsa* may grow through inorganic sulfur disproportionation (Finster et al., 2013; Janssen et al., 1996), and at up to 3.0% relative abundance may play important roles in cycling elemental sulfur, thiosulfate, and sulfite in MIS mats. Sulfur reduction and disproportionation strongly influence photosynthesis and the distribution of cyanobacteria and SOBs in other mat systems (Dick et al., 2018; Klatt et al., 2020; Stal, 2012).

Proteins from other taxa provided additional evidence for seasonal changes in biogeochemistry and metabolism in MIS mats. Diatoms, which likely play important roles in nitrogen cycling within MIS mats (Merz et al., 2021), contributed nearly 16% of peptide spectra in autumn and spring. A carbon fixation protein from another putative phototroph, the purple nonsulfur bacterium *Rhodoferax* (Kaden et al., 2014; Madigan et al., 2000), was highly abundant in summer, suggesting that it may contribute to primary production. Finally, an outer membrane protein from *Methylococcaceae*, a putative methanotroph, was enriched in autumn samples, consistent with more reducing redox conditions within mats in the autumn.

Variation in measured protein relative abundances could reflect both seasonal changes in community composition and changes in protein expression levels within individual taxa. *Phormidium’s* growth during the optimal conditions experienced in summer is reflected in its high relative abundance in both the proteomic and 16S rRNA gene data from those samples. However, in other cases, relative abundances from proteome profiles were not aligned with those from 16S rRNA gene surveys. *Planktothrix* proteins were more often observed in spring than in other seasons, but *Planktothrix* was not the most abundant cyanobacterium in 16S rRNA gene datasets in spring due to the dominance of *Phormidium*. On the other hand, *Planktothrix’*s higher relative abundance of 16S rRNA genes in autumn is more likely a consequence of lower *Phormidium* abundance. In another example, *Beggiatoa* and other sulfur-oxidizing bacteria were present in the bacterial community throughout the year, but the higher abundance of their proteins in autumn likely reflects higher activity and growth.

## Conclusion

This study showed that seasonal changes in irradiance and water chemistry are accompanied by large shifts in microbial community structure and function in the microbial mat of Middle Island Sinkhole. While the current data cannot conclusively disentangle the effects of light versus water chemistry, the proteomic data suggests that light quantity and quality was an important driver of seasonal changes in community composition and metabolism. In addition to changes in the phototrophic community, seasonal shifts in the community of sulfate-reducing bacteria and increased abundance of sulfur-oxidizing *Beggiatoa* in autumn supports the hypothesis that lower light levels lead to an intensified sulfur cycle, perhaps via shifts in photosynthetic rates and/or the balance of oxygenic and anoxygenic photosynthesis. The increased autumn abundance of *Beggiatoa* likely has feedback effects on light available to phototrophs as these sulfur-oxidizing bacteria cover and compete with the cyanobacteria (Biddanda & Weinke, 2020; Klatt et al., 2020; 2021; Klatt, Meyer, et al., 2016). These insights have implications for how seasonality affects the cycling of carbon, sulfur, and oxygen in cyanobacterial mats, and may inform understanding of controls on O_2_ production in widespread redox-stratified mats of the Proterozoic. The results are also relevant to the vulnerability of rare microbial mat ecosystems to environmental change in the modern world; links between microbial composition and function and light and chemistry/hydrology are also likely to manifest in recent anthropogenic environmental change such as changes in water clarity due to invasive dreissenid mussels (Binding et al., 2007) and hydrological shifts associated with climate change, including rapidly fluctuating Great Lakes water levels (Gronewold & Rood, 2019) and changes in groundwater flux.

## Acknowledgements and details of funding

We thank the NOAA Thunder Bay National Marine Sanctuary Dive Unit – John Bright, Russ Green, Phil Hartmeyer, Wayne Lusardi, Stephanie Gandulla, and Tane Casserley – and R/V Storm Ship Captain Travis Smith for field site access, facilities support, and sampling. We also thank Bopaiah Biddanda, Anthony Weinke, Steve Ruberg, Kathryn Rico, Matthew Medina, Judith Klatt, Arjun Chennu, Allen Burton, Michelle Hudson, Lichun Zhang, and Hui Chien Tan for help with field sampling, data collection, processing, and analyses. This work was supported by NSF Grant EAR-1637066 to G.J.D. and University of Michigan Novak Fellowship and Rackham Predoctoral Fellowship to S.L.G.

## Authors contributions

S.L.G. and G.J.D. designed the research. S.L.G., D.G.S., L.E.K.C., and G.J.D. collected samples. S.L.G. and D.G.S. analyzed physicochemical profiles. L.E.K.C. and P.A. performed water chemistry measurements. J.R.W. performed mass spectrometry measurements for proteomics. S.L.G. performed analyses on geochemical, amplicon, and proteomic data sets, with input from N.E.L., J.R.W., and G.J.D. S.L.G. wrote the initial draft of the manuscript. All the authors contributed to the writing of the manuscript.

