## Supplemental Figures and Tables for "Seasonal Shifts in Community Composition and Proteome Expression in a Sulfur-Cycling Cyanobacterial Mat"

**Supplemental Figure 1.** Flat purple mat collected intact with underlying sediment and overlying water column, collected in 2014. The polycarbonate core was 20 cm tall and 7 cm diameter. Purple microbial mat (top half of core tube) was ~mm thick and filaments easily covered tube sides as seen on the right of the image.

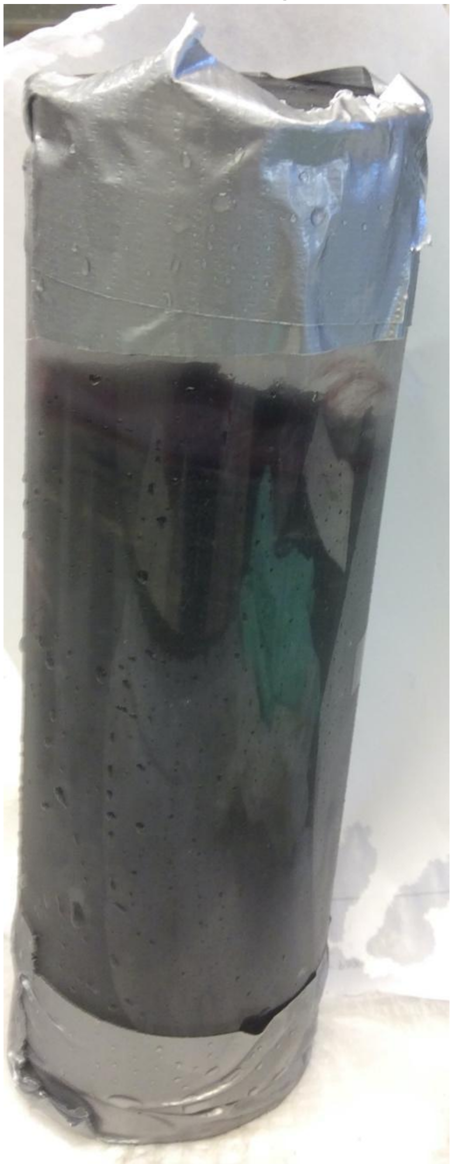

**Supplemental Figure 2.** $\delta^{18}O$ and modeled percentage of groundwater at specific locations in MIS arena in 2016. Samples collected in spring are marked purple, and yellow for samples collected in autumn.

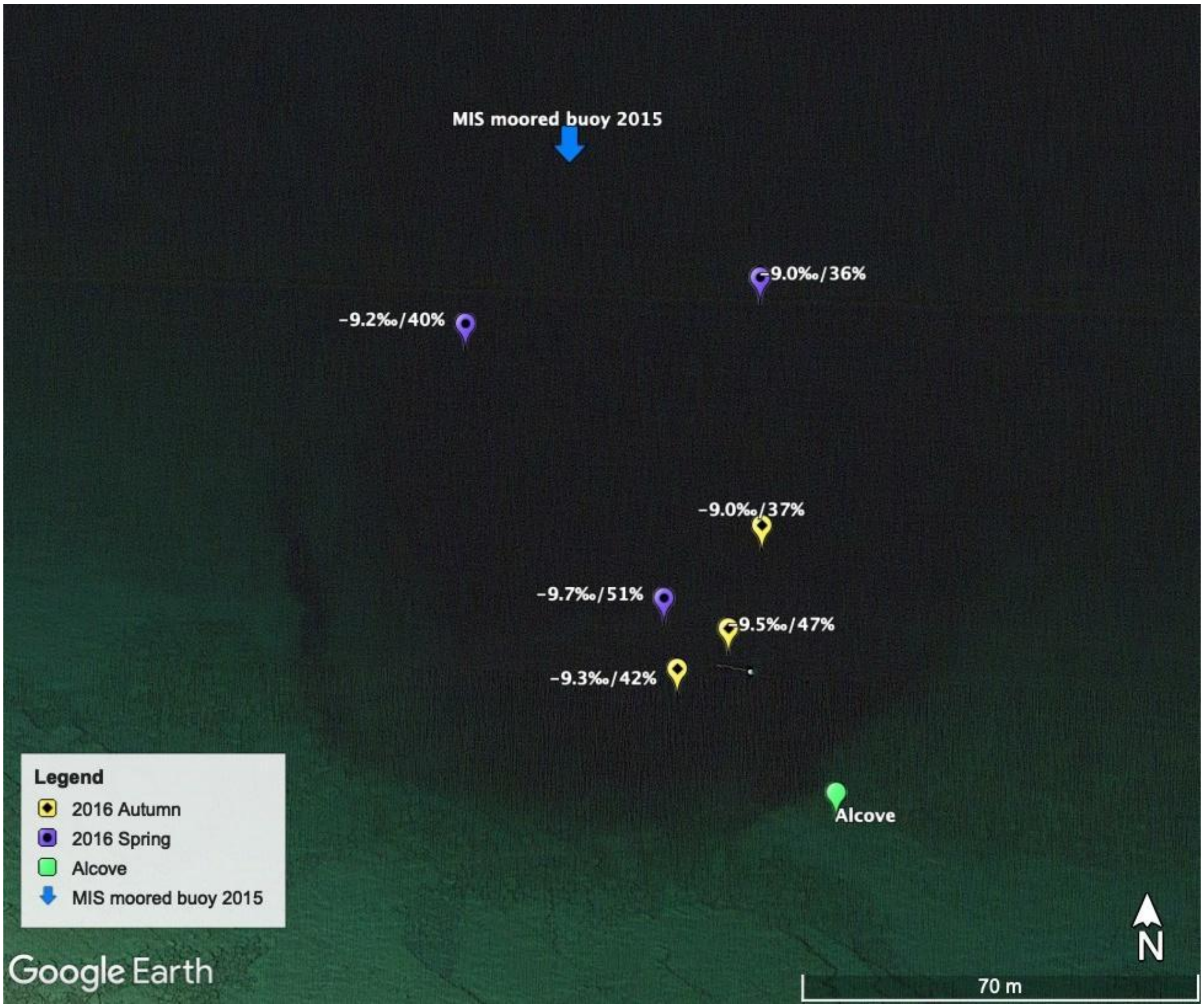

**Supplemental Figure 3.** Quantities of light available at 23 m in the MIS arena or in the open water outside of MIS (“open”) from hyperspectral casts. The available light (y-axis) is plotted for each wavelength (x-axis) of PAR. Different sampling days are represented with colors, and separate casts from the same day are in solid or dashed lines.

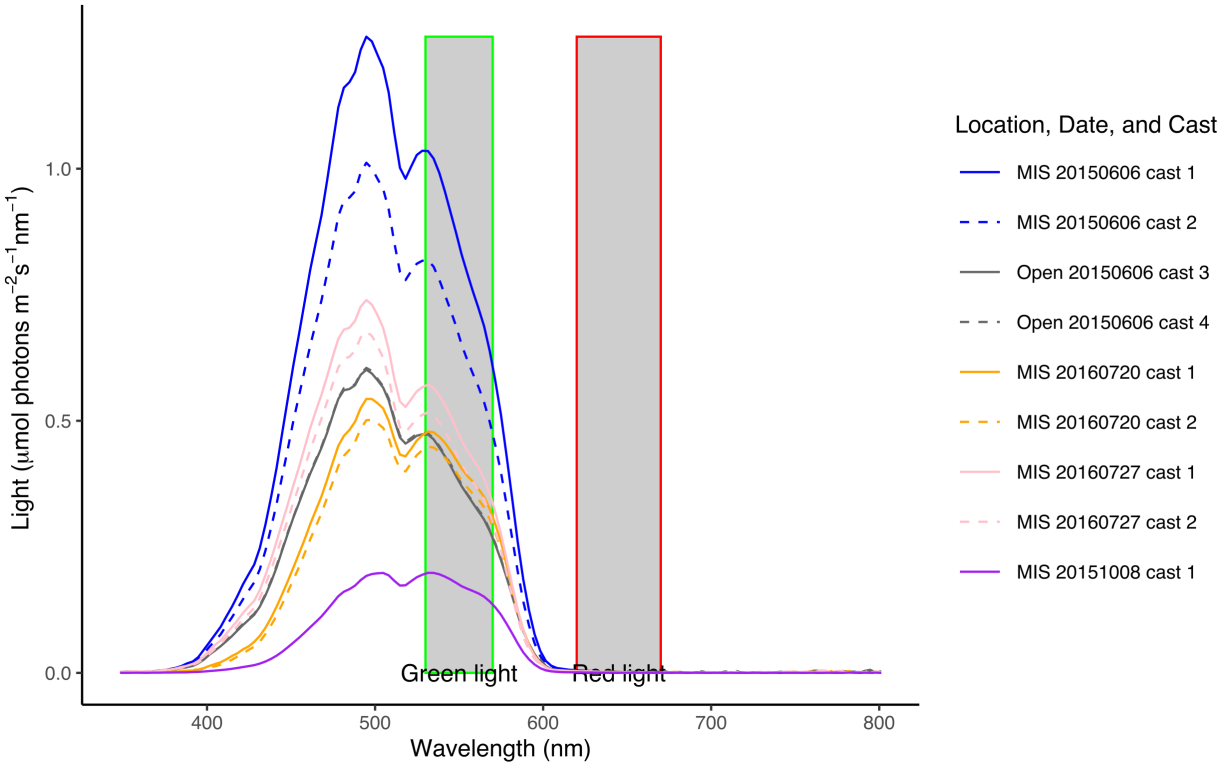

**Supplemental Figure 4.** Calculated *k*-extinction coefficients at 23 m in the MIS arena or away from the arena (“open”) from hyperspectral casts. The extinction coefficient (y-axis) is plotted for each wavelength (x-axis) of PAR. Different sampling days are represented with colors, and separate casts from the same day are in solid or dashed lines.

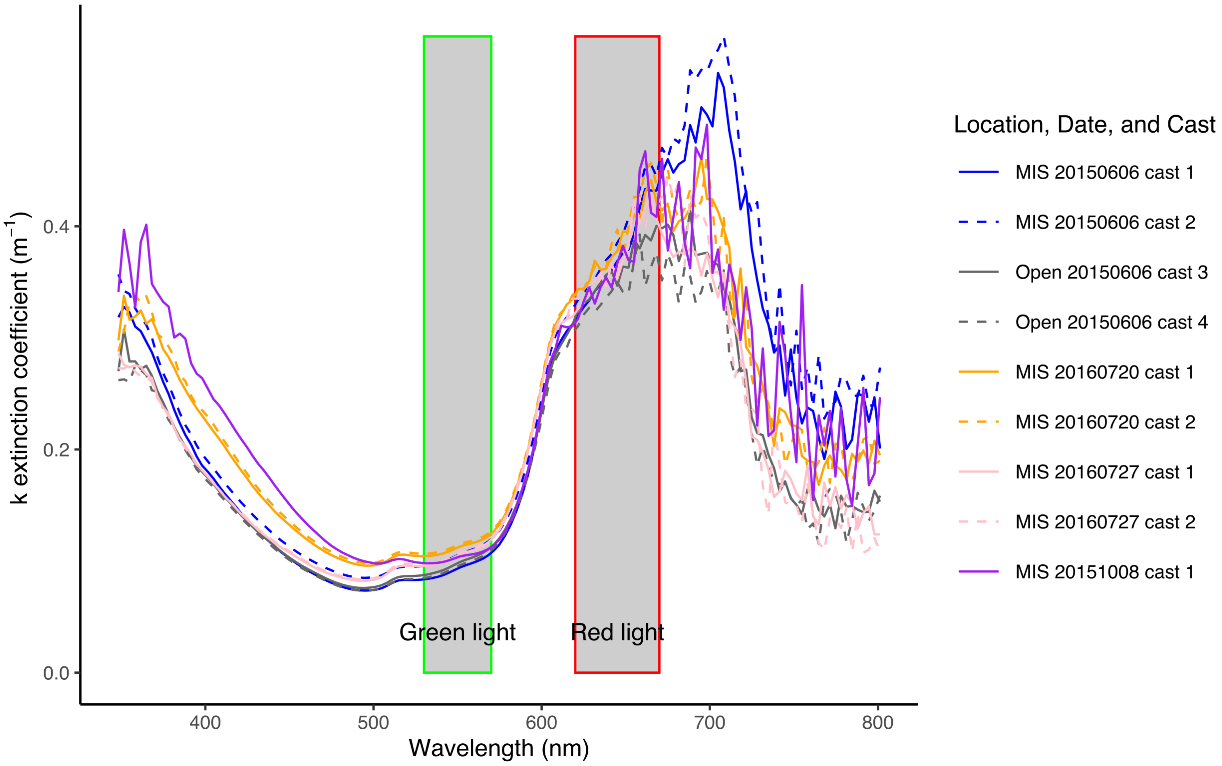

**Supplemental Figure 5.** Specific conductivity measured with hand-held probe or calculated from ion chemistry (IC and ICPMS) in water samples. Units are µS $cm^{-1}$.

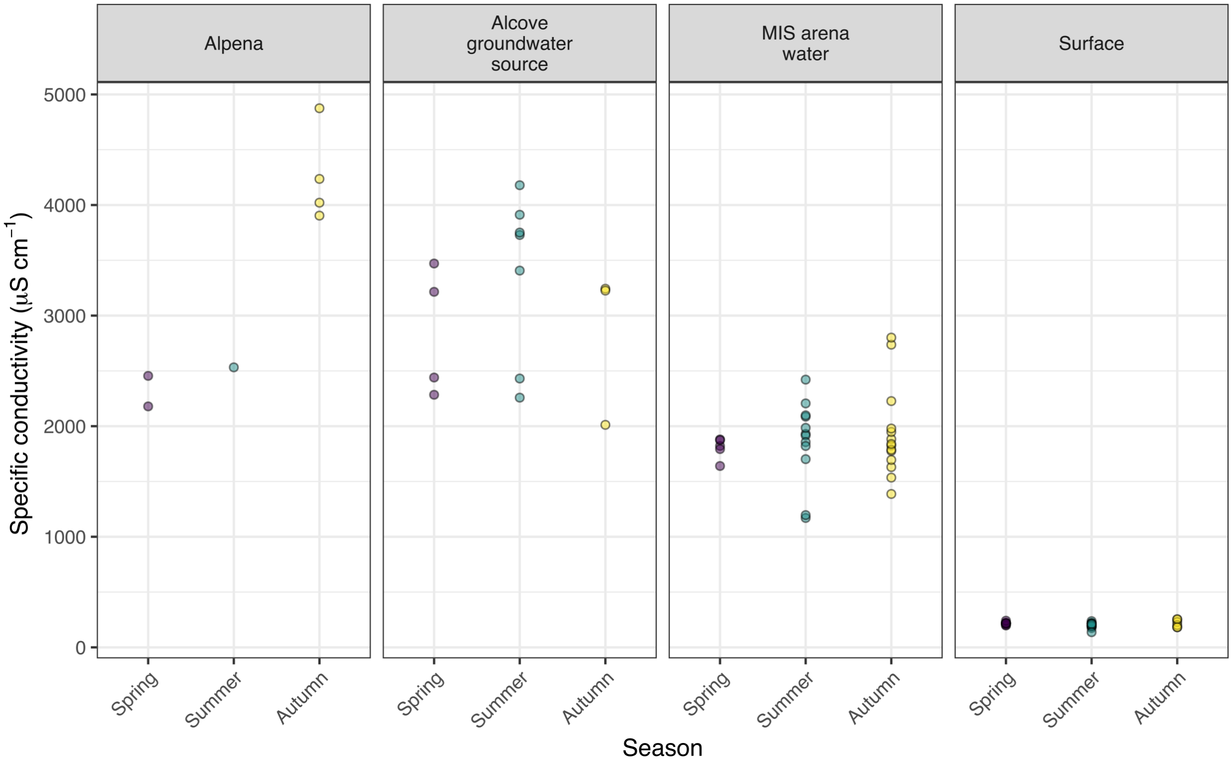

**Supplemental Figure 6.** *d-excess* (difference between measured values of δD and modeled δD from measured values of $\delta^{18}O$) for water samples at each location per season. The mean for each season is presented as the colored horizontal line, and box and whiskers summarize 25-75th percentiles of observations (colored circles). For context, the average *d-excess* for the Great Lakes is +3.2‰.

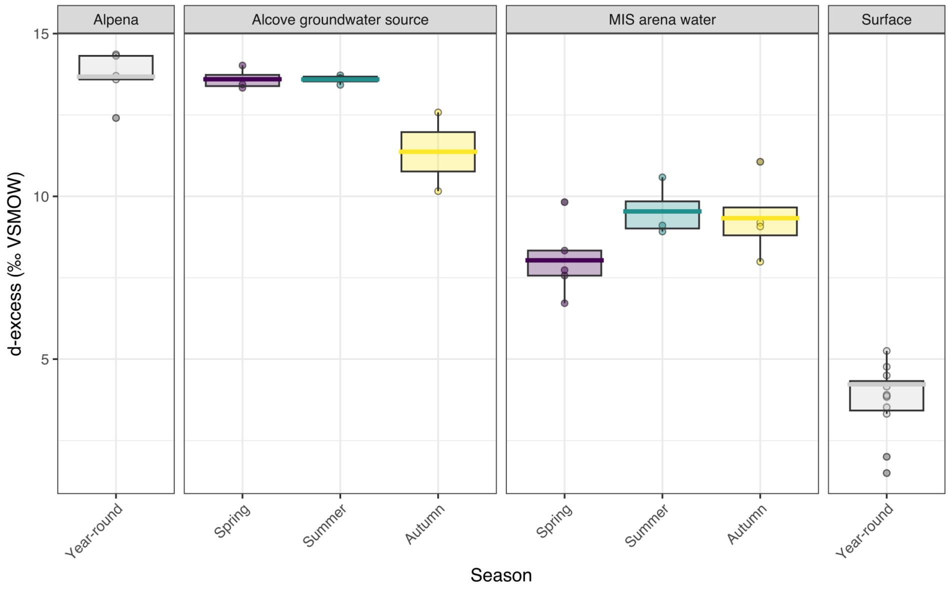

**Supplemental Figure 7.** Measured ion concentrations in water samples compared to predicted ion concentrations based on δ$^{18}$O-derived linear mixing model. Colored points represent measurements of ions of interest: A, sulfate; B, calcium; C, fluoride; D, sodium; E, magnesium, and F, chloride. The horizontal colored line is the mean measured concentration per season unit. For each season, the gray box represents the mean predicted concentrations and standard deviations using the concentrations measured in Alpena fountain and the isotope-derived linear mixing model between surface water and Alpena fountain water.

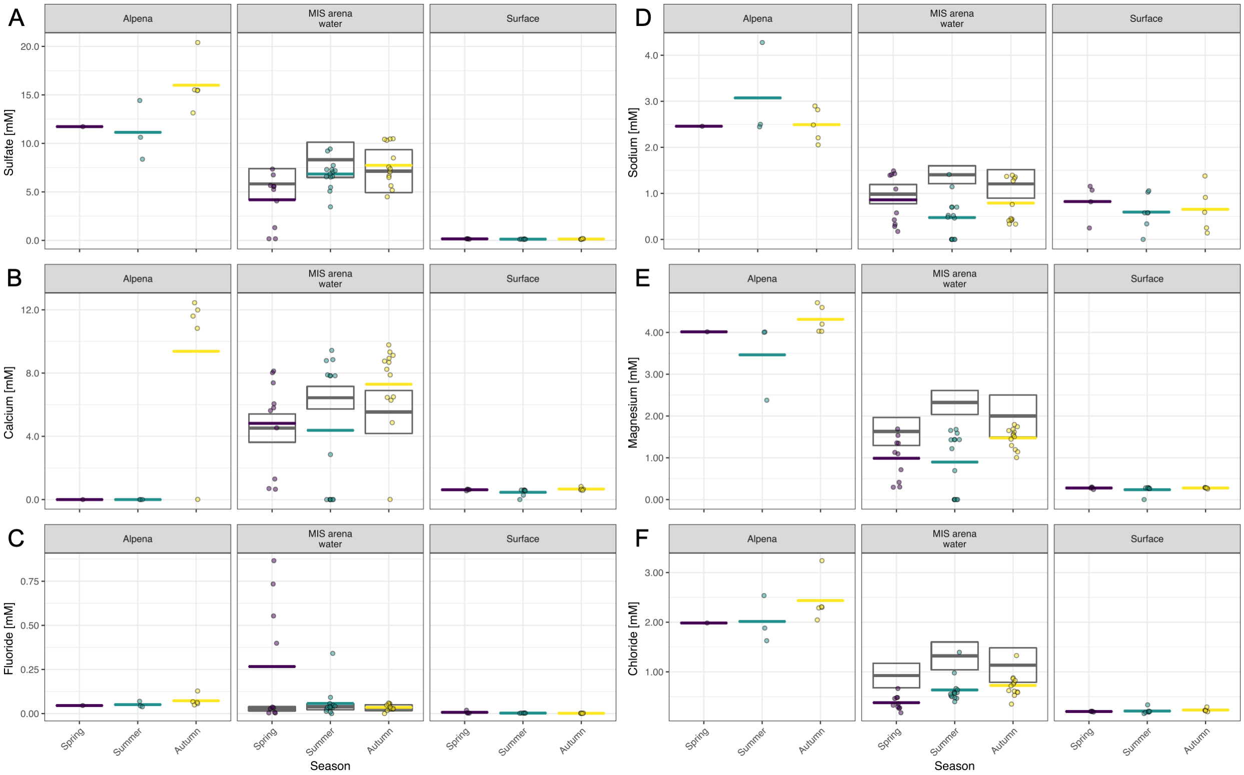

**Supplemental Figure 8.** Relative abundance of relevant bacterial taxonomic groups in samples, grouped by month and year. Relative abundances of key genera were summed and presented, and classes and phyla without those genera are represented.

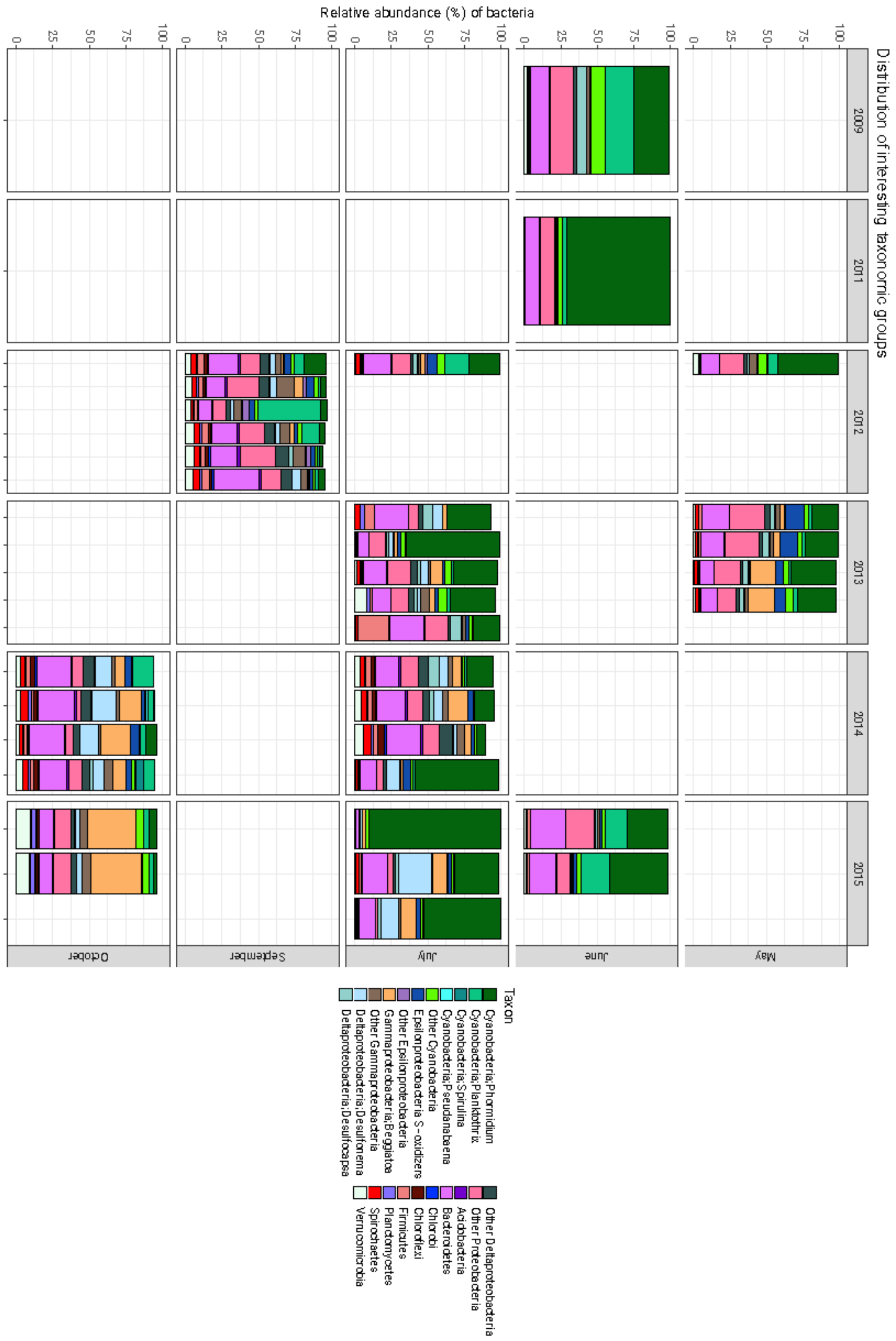

**Supplemental Figure 9.** Correlation network showing significant ($p < 0.001$) relationships in relative abundances of 60 genera. Bray-Curtis distance matrix was calculated from read counts of genera and used to generate a correlation network.

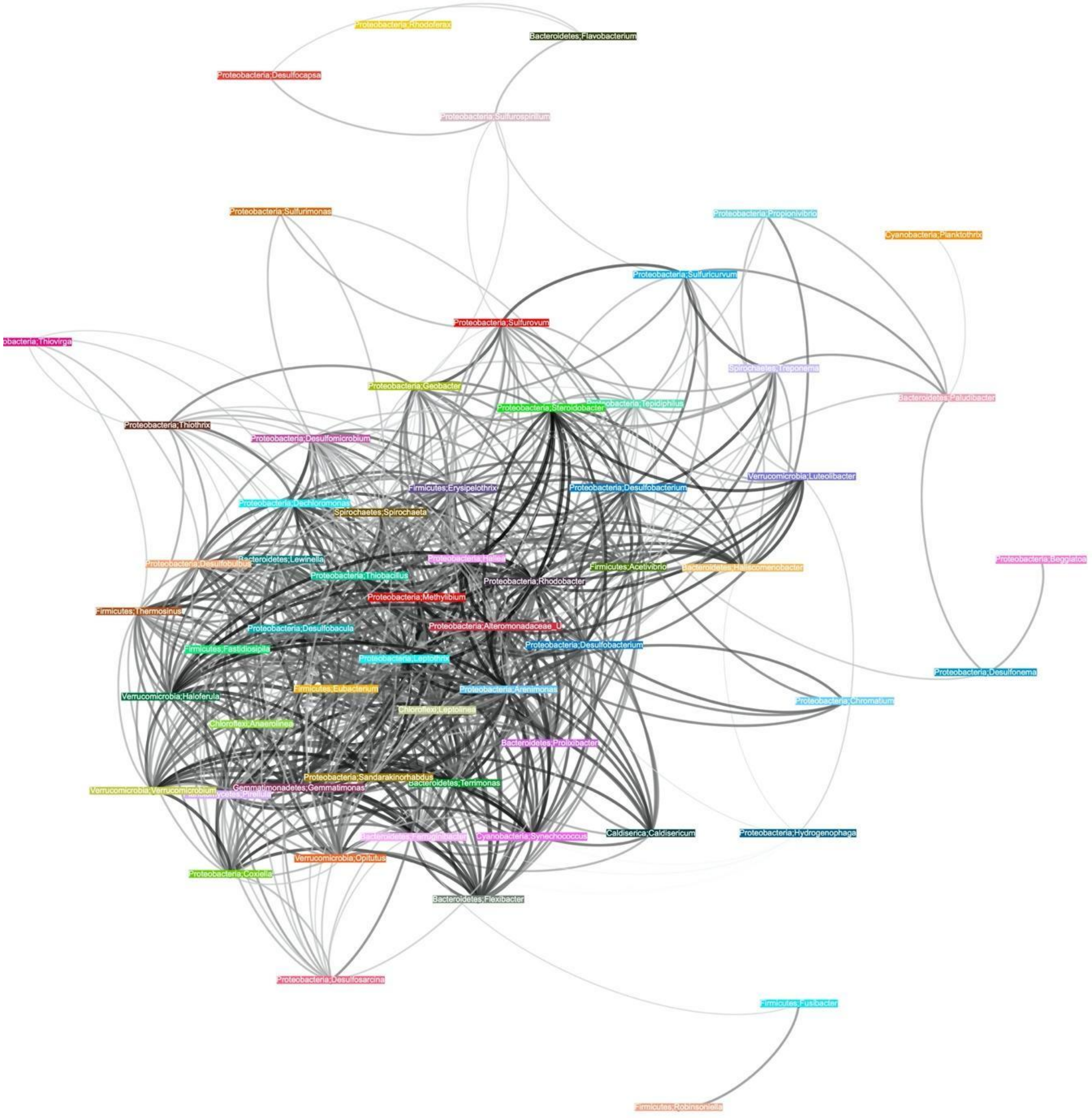

**Supplemental Table 1**. (this and next 4 pages) Summary table of metagenome- assembled-genomic (MAG) bins from MIS metagenomic co-assembly of 15 samples. Our oscillatorial cyanobacterial bins had 95.8% completion/11.3% redundancy (*Pseudanabaena*, 4.3Mbp), 70.5%/56.3% (*Phormidium*, 11.0Mbp), 91.6%/42.3% (*Planktothrix*, 5.7Mbp), 22.5%/1.41% (*Spirulina*, 1.12Mbp) as well as 527kbp of cyanobacterial scaffolds that were not able to be confidently binned.

| Bin name | Classification | Bin Quality | Length (bp) | Completion (%) | Redundancy (%) | Mean gDNA coverage | Mean cDNA coverage |
|---|---|---|---|---|---|---|---|
| bin_208 | Alphaproteobacteria; Rickettsia | Medium | 1036760 | 71.83 | 0.0 | 0.04 | 0.0 |
| bin_118 | Bacteria | High | 3917186 | 98.59 | 1.41 | 0.77 | 3.62 |
| bin_156 | Bacteria | Low | 3085822 | 63.38 | 0.0 | 1.20 | 0.13 |
| bin_146 | Bacteria | Low | 2304693 | 0.0 | 0.0 | 0.14 | 0.02 |
| bin_51 | Bacteria | Low | 1580267 | 0.0 | 0.0 | 0.55 | 0.16 |
| bin_220 | Bacteria | Low | 1483107 | 19.72 | 0.0 | 0.68 | 0.03 |
| bin_103 | Bacteria | Low | 1266519 | 0.0 | 0.0 | 0.57 | 0.02 |
| bin_157 | Bacteria | Low | 163623 | 0.0 | 0.0 | 0.05 | 0.0 |
| bin_168 | Bacteria | Low | 157396 | 0.0 | 0.0 | 0.04 | 0.0 |
| bin_182 | Bacteria | Medium | 2968070 | 88.73 | 1.41 | 0.06 | 3.84 |
| bin_50 | Bacteria | Medium | 3858105 | 87.32 | 8.45 | 1.06 | 0.39 |
| Bin_7_3-contigs | Bacteria | Medium | 3971796 | 85.92 | 1.41 | 5.29 | 0.54 |
| bin_212 | Bacteria | Medium | 2393843 | 83.10 | 5.63 | 1.18 | 0.77 |
| bin_47 | Bacteria | Medium | 3407323 | 53.52 | 1.41 | 0.19 | 0.02 |
| bin_105 | Bacteria | Medium | 4024241 | 70.42 | 1.41 | 4.10 | 0.05 |
| bin_64 | Bacteria; Silvanigrellales | Low | 232602 | 0.0 | 0.0 | 0.02 | 0.04 |
| bin_250 | Bacteria; Vampirococcus | Medium | 1008733 | 53.52 | 1.41 | 0.15 | 0.02 |
| bin_9 | Bacteroidetes | High | 4643096 | 100.0 | 2.82 | 0.76 | 0.04 |
| bin_33 | Bacteroidetes | High | 3200533 | 98.59 | 2.82 | 0.33 | 0.37 |
| bin_28 | Bacteroidetes | High | 4812335 | 98.59 | 1.41 | 1.22 | 3.04 |
| bin_207 | Bacteroidetes | High | 2366644 | 92.96 | 0.0 | 0.31 | 3.83 |
| bin_228 | Bacteroidetes | High | 6581157 | 94.37 | 4.23 | 0.14 | 0.01 |
| bin_57 | Bacteroidetes | High | 3922016 | 97.18 | 1.41 | 0.23 | 3.00 |
| bin_160 | Bacteroidetes | Low | 1482139 | 43.66 | 0.0 | 0.76 | 0.07 |
| bin_202 | Bacteroidetes | Low | 3626904 | 32.39 | 2.82 | 0.65 | 0.12 |
| bin_87 | Bacteroidetes | Low | 720606 | 42.25 | 0.0 | 1.20 | 4.46 |
| bin_124 | Bacteroidetes | Low | 1128359 | 36.62 | 0.0 | 0.08 | 0.02 |
| bin_224 | Bacteroidetes | Low | 1116623 | 35.21 | 0.0 | 0.38 | 0.03 |
| bin_113 | Bacteroidetes | Low | 823305 | 0.0 | 0.0 | 0.26 | 0.03 |
| bin_23 | Bacteroidetes | Low | 347815 | 0.0 | 0.0 | 0.91 | 0.03 |
| bin_261 | Bacteroidetes | Low | 54079 | 0.0 | 0.0 | 0.07 | 0.0 |
| bin_178 | Bacteroidetes | Low | 371388 | 0.0 | 0.0 | 0.25 | 0.04 |
| bin_92 | Bacteroidetes | Medium | 3783221 | 87.32 | 1.41 | 0.40 | 0.04 |
| bin_69 | Bacteroidetes | Medium | 3315494 | 91.55 | 4.23 | 0.69 | 0.04 |

**Supplemental Table 1**.

| | | | | | | | |
|---|---|---|---|---|---|---|---|
| bin_88 | Bacteroidetes | Medium | 2317404 | 80.28 | 2.82 | 0.36 | 0.04 |
| bin_153 | Bacteroidetes | Medium | 2962889 | 71.83 | 8.45 | 0.17 | 39.29 |
| bin_117 | Bacteroidetes | Medium | 1688616 | 63.38 | 0.0 | 1.43 | 0.06 |
| bin_239 | Bacteroidetes | Medium | 2545173 | 69.01 | 1.41 | 0.29 | 0.03 |
| bin_3 | Bacteroidetes | Medium | 4300737 | 61.97 | 1.41 | 0.36 | 0.02 |
| bin_54 | Bacteroidetes; Bernardetia | Low | 571641 | 47.89 | 1.41 | 0.76 | 84.01 |
| bin_176 | Bacteroidetes; Draconibacterium | Low | 945589 | 0.0 | 0.0 | 0.66 | 0.09 |
| bin_197 | Bacteroidetes; Draconibacterium | Low | 1178942 | 0.0 | 0.0 | 0.33 | 0.09 |
| bin_169 | Bacteroidetes; Draconibacterium | Medium | 2672817 | 54.93 | 0.0 | 0.49 | 0.16 |
| bin_242 | Bacteroidetes; Flavobacterium | Low | 788631 | 0.0 | 0.0 | 0.31 | 0.03 |
| bin_96 | Bacteroidetes; Flavobacterium | Low | 339985 | 0.0 | 0.0 | 9.98 | 0.24 |
| bin_227 | Bacteroidetes; Flavobacterium | Medium | 2930134 | 76.06 | 0.0 | 0.94 | 0.04 |
| bin_75 | Bacteroidetes; Paludibacter | Low | 602309 | 46.48 | 0.0 | 2.13 | 0.06 |
| bin_100 | Bacteroidetes; Paludibacter | Low | 1400511 | 0.0 | 0.0 | 4.63 | 0.09 |
| bin_132 | Bacteroidetes; Paludibacter | Medium | 4302557 | 84.51 | 1.41 | 1.57 | 2.41 |
| bin_112 | Bacteroidetes; Paludibacter | Medium | 2800085 | 64.79 | 4.23 | 3.58 | 0.11 |
| bin_200 | Betaproteobacteria | Low | 218076 | 28.17 | 0.0 | 1.73 | 0.05 |
| bin_111 | Betaproteobacteria | Low | 637507 | 0.0 | 0.0 | 1.85 | 0.04 |
| bin_127 | Betaproteobacteria | Medium | 3724154 | 70.42 | 2.82 | 1.17 | 0.05 |
| bin_91 | Betaproteobacteria | Medium | 3172373 | 80.28 | 0.0 | 1.99 | 0.06 |
| bin_151 | Betaproteobacteria | Medium | 2184168 | 76.06 | 1.41 | 1.82 | 0.05 |
| bin_77 | Betaproteobacteria; Candidatus Accumulibacter | Low | 427799 | 0.0 | 0.0 | 1.05 | 0.0 |
| bin_122 | Betaproteobacteria; Candidatus Accumulibacter | Medium | 3455847 | 74.65 | 2.82 | 0.79 | 0.01 |
| bin_131 | Betaproteobacteria; Collimonas | High | 1613140 | 97.18 | 0.0 | 0.03 | 0.01 |
| Bin_4_4-contigs | Betaproteobacteria; Hydrogenophaga | High | 3519382 | 92.96 | 1.41 | 10.93 | 0.24 |
| bin_70 | Betaproteobacteria; Hydrogenophaga | Medium | 2249573 | 70.42 | 0.0 | 4.68 | 0.11 |
| Bin_7_6-contigs | Betaproteobacteria; Methylibium | Low | 1954091 | 29.58 | 0.0 | 1.38 | 0.02 |
| Bin_4_1-contigs | Betaproteobacteria; Rhodoferax | n/a | 4187395 | 90.14 | 90.14 | 7.22 | 1.56 |
| bin_44 | Betaproteobacteria; Rhodoferax | Low | 2317582 | 45.07 | 0.0 | 1.30 | 0.14 |
| bin_106 | Betaproteobacteria; Rhodoferax | Low | 2418005 | 0.0 | 0.0 | 3.78 | 0.08 |

| Supplemental Table 1. | | | | | | | |
|---|---|---|---|---|---|---|---|
| bin_114 | Betaproteobacteria; Rhodoferax | Low | 1033036 | 0.0 | 0.0 | 0.72 | 0.02 |
| bin_38 | Betaproteobacteria; Rhodoferax | Low | 1189669 | 0.0 | 0.0 | 2.36 | 0.05 |
| bin_167 | Betaproteobacteria; Rhodoferax | Low | 569153 | 0.0 | 0.0 | 1.39 | 0.22 |
| bin_229 | Betaproteobacteria; Rhodoferax | Medium | 4595121 | 87.32 | 1.41 | 0.48 | 0.26 |
| bin_246 | Betaproteobacteria; Rhodoferax | Medium | 2662737 | 69.01 | 2.82 | 1.46 | 0.04 |
| bin_46 | Betaproteobacteria; Rhodoferax | Medium | 2729705 | 66.20 | 5.63 | 2.00 | 0.05 |
| bin_144 | Betaproteobacteria; Sideroxydans | Low | 1077631 | 38.03 | 4.23 | 0.43 | 0.61 |
| Bin_4_3_4-contigs | Betaproteobacteria; Sulfuritalea | Medium | 3046083 | 67.61 | 1.41 | 0.22 | 0.65 |
| bin_193 | Chlorobi | Medium | 2240036 | 67.61 | 1.41 | 0.11 | 0.89 |
| bin_123 | Chloroflexi | Low | 2116432 | 38.03 | 2.82 | 1.22 | 0.03 |
| bin_120 | Chloroflexi | Medium | 2823555 | 85.92 | 2.82 | 1.02 | 2.73 |
| bin_187 | Chloroflexi | Medium | 4444501 | 61.97 | 1.41 | 0.43 | 35.36 |
| bin_154 | Chloroflexi; Anaerolinea | Low | 820620 | 22.54 | 1.41 | 0.73 | 0.05 |
| bin_179 | Chloroflexi; Anaerolinea | Low | 919484 | 0.0 | 0.0 | 0.79 | 0.04 |
| bin_190 | Chloroflexi; Anaerolinea | Low | 406092 | 0.0 | 0.0 | 0.53 | 0.04 |
| bin_232 | Chloroflexi; Anaerolinea | Low | 476289 | 0.0 | 0.0 | 0.16 | 0.0 |
| bin_8 | Chloroflexi; Anaerolinea | Medium | 4061672 | 63.38 | 1.41 | 0.34 | 0.01 |
| bin_141 | Chloroflexi; Anaerolinea | Medium | 3352447 | 49.30 | 1.41 | 0.50 | 0.02 |
| bin_43 | Chloroflexi; Anaerolinea | Medium | 1290979 | 56.34 | 1.41 | 0.22 | 0.06 |
| bin_180 | Cyanobacteria; Oscillatoriales | Low | 527123 | 0.0 | 0.0 | 0.73 | 0.05 |
| Bin_1 | Cyanobacteria; Phormidium | n/a | 11033395 | 70.42 | 56.34 | 203.01 | 181.21 |
| bin_235_243 | Cyanobacteria; Planktothrix | n/a | 5709010 | 91.55 | 42.25 | 28.57 | 1.54 |
| bin_143 | Cyanobacteria; Pseudanabaena | Medium | 4277160 | 95.77 | 11.27 | 0.46 | 0.07 |
| bin_256 | Cyanobacteria; Spirulina | Low | 1128073 | 22.54 | 1.41 | 1.18 | 0.98 |
| bin_90 | Deltaproteobacteria | Low | 1629916 | 43.66 | 0.0 | 0.79 | 0.03 |
| bin_145_29 | Deltaproteobacteria | Low | 998016 | 0.0 | 0.0 | 0.13 | 0.02 |
| bin_5 | Deltaproteobacteria | Medium | 2954443 | 77.46 | 0.0 | 0.34 | 0.02 |
| bin_65 | Deltaproteobacteria; Bdellovibrio | Low | 434310 | 0.0 | 0.0 | 0.07 | 0.0 |
| bin_204 | Deltaproteobacteria; Desulfobacteraceae | Low | 1256801 | 45.07 | 1.41 | 6.42 | 0.21 |

| Supplemental Table 1. | | | | | | | |
|---|---|---|---|---|---|---|---|
| bin_101 | Deltaproteobacteria; Desulfobacteraceae | Medium | 3961593 | 88.73 | 4.23 | 0.41 | 0.03 |
| bin_83 | Deltaproteobacteria; Desulfobacteraceae | Medium | 2395034 | 67.61 | 4.23 | 0.95 | 2.37 |
| bin_26 | Deltaproteobacteria; Desulfobacula | High | 3688401 | 97.18 | 4.23 | 0.20 | 0.03 |
| bin_102 | Deltaproteobacteria; Desulfobulbaceae | Medium | 3062052 | 71.83 | 1.41 | 0.57 | 0.03 |
| bin_170 | Deltaproteobacteria; Desulfobulbaceae | Medium | 2433296 | 61.97 | 5.63 | 0.70 | 0.03 |
| bin_61 | Deltaproteobacteria; Desulfocapsa | Low | 1160311 | 21.13 | 0.0 | 1.41 | 0.07 |
| bin_85 | Deltaproteobacteria; Desulfocapsa | Low | 574286 | 0.0 | 0.0 | 2.22 | 15.58 |
| bin_6 | Deltaproteobacteria; Desulfomicrobium | Low | 1947632 | 26.76 | 0.0 | 1.24 | 1.38 |
| bin_209 | Deltaproteobacteria; Halobacteriovorax | High | 3253134 | 91.55 | 2.82 | 0.09 | 0.77 |
| bin_67 | Epsilonproteobacteria | Low | 665089 | 21.13 | 0.0 | 1.23 | 63.48 |
| bin_25 | Epsilonproteobacteria | Low | 317211 | 30.99 | 0.0 | 0.46 | 0.15 |
| Bin_6_9-contigs | Epsilonproteobacteria; Arcobacter | High | 1997449 | 97.18 | 0.0 | 7.00 | 0.11 |
| bin_49 | Epsilonproteobacteria; Sulfuricurvum | High | 2217859 | 95.77 | 0.0 | 1.37 | 0.12 |
| Bin_6_3-contigs | Epsilonproteobacteria; Sulfurimonas | High | 2205377 | 98.59 | 1.41 | 0.42 | 0.02 |
| bin_94 | Epsilonproteobacteria; Sulfurospirillum | Low | 501389 | 0.0 | 0.0 | 1.07 | 0.07 |
| bin_233 | Epsilonproteobacteria; Sulfurospirillum | Medium | 797513 | 53.52 | 0.0 | 2.24 | 0.14 |
| Bin_10_2-contigs | Eukaryota | Low | 8793166 | 32.53 | 3.61 | 0.02 | 0.01 |
| bin_2 | Eukaryota | n/a | 13858755 | 21.69 | 25.30 | 0.03 | 0.37 |
| bin_159 | Fibrobacteres | Low | 900344 | 0.0 | 0.0 | 0.60 | 0.03 |
| bin_194 | Firmicutes - Clostridia | Low | 329682 | 29.58 | 1.41 | 0.26 | 0.0 |
| bin_86 | Firmicutes - Clostridia | Medium | 3921580 | 60.56 | 1.41 | 0.26 | 0.03 |
| bin_78 | Gammaproteobacteria | Low | 3197050 | 46.48 | 5.63 | 1.01 | 0.04 |
| bin_73 | Gammaproteobacteria | Low | 2387643 | 33.80 | 4.23 | 1.64 | 0.09 |
| bin_164 | Gammaproteobacteria | Low | 948971 | 0.0 | 0.0 | 2.04 | 0.30 |
| bin_37 | Gammaproteobacteria | Low | 595635 | 0.0 | 0.0 | 5.24 | 0.26 |
| bin_99 | Gammaproteobacteria | Low | 540621 | 0.0 | 0.0 | 0.43 | 0.0 |
| bin_45 | Gammaproteobacteria; Methylococcaceae | High | 2731051 | 95.77 | 2.82 | 0.29 | 0.10 |
| bin_149 | Gammaproteobacteria; Methylovulum | Low | 1412910 | 56.34 | 4.23 | 0.05 | 4.96 |
| bin_234 | Gammaproteobacteria; Thioploca | Low | 750004 | 0.0 | 0.0 | 0.13 | 0.0 |
| bin_35 | Gammaproteobacteria; Thioploca | Low | 666513 | 0.0 | 0.0 | 13.76 | 0.22 |
| bin_247 | Gammaproteobacteria; Thioploca | Low | 902565 | 0.0 | 0.0 | 2.75 | 0.07 |

| | | | | | | | |
|---|---|---|---|---|---|---|---|
| bin_188 | Gammaproteobacteria; Thioploca | Low | 687171 | 0.0 | 0.0 | 2.99 | 0.10 |
| bin_158 | Gammaproteobacteria; Thiotrichaceae | Medium | 2018828 | 77.46 | 1.41 | 0.24 | 13.05 |
| bin_16 | Spirochaetes | Low | 1665686 | 0.0 | 0.0 | 0.81 | 0.03 |
| bin_134 | Spirochaetes | Low | 1818317 | 23.94 | 2.82 | 0.29 | 0.03 |
| bin_162 | Spirochaetes | Low | 1282009 | 21.13 | 4.23 | 0.87 | 0.04 |
| bin_219 | Spirochaetes | Low | 976721 | 0.0 | 0.0 | 0.24 | 0.02 |
| bin_172 | Spirochaetes | Low | 1252337 | 0.0 | 0.0 | 0.86 | 0.03 |
| bin_201 | Spirochaetes | Low | 437961 | 0.0 | 0.0 | 0.41 | 0.03 |
| bin_257 | Spirochaetes | Low | 194932 | 0.0 | 0.0 | 0.33 | 0.03 |
| bin_199 | Spirochaetes | Medium | 3969302 | 90.14 | 7.04 | 1.31 | 0.73 |
| bin_251 | Spirochaetes | Medium | 2806768 | 43.66 | 1.41 | 1.10 | 0.03 |
| bin_262 | Tenericutes | Low | 629939 | 46.48 | 1.41 | 0.09 | 0.0 |
| bin_79 | Unknown | Low | 575930 | 40.85 | 0.0 | 0.04 | 2.68 |
| Bin_3_1 | Unknown | Low | 23570043 | 87.32 | 254.93 | 0.31 | 20.59 |
| Bin_3_3 | Unknown | Low | 19802335 | 90.14 | 236.62 | 6.19 | 17.77 |
| bin_155 | Verrucomicrobia | Low | 548848 | 50.70 | 1.41 | 0.51 | 0.03 |
| bin_66 | Verrucomicrobia | Low | 2006576 | 50.70 | 0.0 | 0.13 | 0.01 |
| bin_72 | Verrucomicrobia | Low | 1332738 | 45.07 | 0.0 | 0.42 | 0.05 |
| bin_165 | Verrucomicrobia | Low | 450480 | 0.0 | 0.0 | 0.92 | 0.03 |
| bin_89 | Verrucomicrobia; Akkermansia | Medium | 3062395 | 70.42 | 2.82 | 0.39 | 5.75 |

**Supplemental Table 2.** Light quality and quantity metrics from hyperspectral profiles. Seven hyperspectral casts in MIS arena and 2 open water casts were conducted in 2015-2016, from which we determined *k*-extinction coefficients and available energy (µmol photons $m^{-2}$ $s^{-1}$ $nm^{-1}$) for each measured wavelength of light between 350-800 nm. This summarizes the *k*-extinction coefficient and energy at depth for the range of green wavelengths (530-570 nm) and the range of red wavelengths (620-670 nm), per cast, per day, and per month.

| | | **2015 0606 cast 1** | **2015 0606 cast 2** | **2015 0606 cast 3** | **2015 0606 cast 4** | **2015 1008 cast 1** | **2016 0720 cast 1** | **2016 0720 cast 2** | **2016 0727 cast 1** | **2016 0727 cast 2** |
|---|---|---|---|---|---|---|---|---|---|---|
| **k-extinction coefficients ($m^{-1}$)** | **Green** | 0.093 | 0.105 | 0.098 | 0.095 | 0.103 | 0.112 | 0.114 | 0.105 | 0.107 |
| | **Red** | 0.374 | 0.391 | 0.361 | 0.345 | 0.373 | 0.385 | 0.392 | 0.374 | 0.373 |
| **energy at 23 m (µmol photons $nm^{-1}$)** | **Green** | 0.852 | 0.669 | 0.382 | 0.385 | 0.175 | 0.413 | 0.388 | 0.474 | 0.426 |
| | **Red** | 0.002 | 0.001 | 0.001 | 0.002 | 0.000 | 0.001 | 0.001 | 0.002 | 0.001 |
| **average daily k-extinction coefficient ($m^{-1}$)** | **Green** | 0.099 | 0.099 | 0.097 | 0.097 | NA | 0.113 | 0.113 | 0.106 | 0.106 |
| | **Red** | 0.383 | 0.383 | 0.353 | 0.353 | NA | 0.389 | 0.389 | 0.373 | 0.373 |
| **standard deviation** | **Green** | 0.009 | 0.009 | 0.007 | 0.007 | NA | 0.006 | 0.006 | 0.007 | 0.007 |
| | **Red** | 0.041 | 0.041 | 0.024 | 0.024 | NA | 0.035 | 0.035 | 0.032 | 0.032 |
| **average monthly k-extinction coefficient ($m^{-1}$)** | **Green** | NA | NA | NA | NA | NA | 0.109 | 0.109 | 0.109 | 0.109 |
| | **Red** | NA | NA | NA | NA | NA | 0.381 | 0.381 | 0.381 | 0.381 |
| **standard deviation** | **Green** | NA | NA | NA | NA | NA | 0.007 | 0.007 | 0.007 | 0.007 |
| | **Red** | NA | NA | NA | NA | NA | 0.034 | 0.034 | 0.034 | 0.034 |

**Supplemental Table 3.** Specific conductivity in water parcels. Units are in µS $cm^{-1}$.

| Date | Season | Alpena | Alcove | Arena | Surface |
|---|---|---|---|---|---|
| 20120531 | Spring | | | 1821 | 219 |
| 20120725 | Summer | | | 1930 | 217 |
| 20120927 | Autumn | 4875 | | 1883 | 251 |
| 20130509 | Spring | | | 1793 | 220 |
| 20130718 | Summer | | 3407 | 2086 | 206 |
| 20130928 | Autumn | | | 1825 | |
| 20140724 | Summer | | 3729 | 1702 | 175 |
| 20140924 | Autumn | 3904 | 3243 | 1630 | 182 |
| 20150604 | Spring | 2179 | 2440 | | 203 |
| 20150604 | Spring | | 2284 | | 214 |
| 20150604 | Spring | | | | 218 |
| 20150731 | Summer | | 2431 | 1916 | 237 |
| 20150731 | Summer | | 3912 | 2206 | 194 |
| 20150731 | Summer | | 4179 | 2098 | 195 |
| 20150731 | Summer | | | | |
| 20151007 | Autumn | | 2012 | 1948 | 229 |
| 20151007 | Autumn | 4021 | | 2737 | 201 |
| 20151007 | Autumn | 4236 | | 2801 | |
| 20160609 | Spring | | 3215 | 1880 | 219 |
| 20160609 | Spring | | | 1640 | |
| 20160722 | Summer | | | 1170 | 217 |
| 20160928 | Autumn | | | 1535 | |
| 20160928 | Autumn | | | 2227 | |
| 20160928 | Autumn | | | 1979 | 255 |
| 20160928 | Autumn | | 3226 | 1776 | 182 |
| 20170601 | Spring | | 3471 | | 198 |
| 20170601 | Spring | | | | 241 |
| 20170809 | Summer | | 3751 | | |
| 20170809 | Summer | | 2258 | 1856 | 218 |
| 20170809 | Summer | | | | 192 |
| 20170810 | Summer | | | 2421 | 192 |
| 20170810 | Summer | | | 1821 | 205 |
| 20170910 | Autumn | | | 1787 | |
| 20170926 | Autumn | | | 1839 | |
| 20170927 | Autumn | | | 1695 | |
| 20170928 | Autumn | | | 1387 | |
| 20180710 | Summer | | | 1985 | 211 |
| 20180710 | Summer | 2532 | | | 137 |
| 20190612 | Spring | 2455 | | | 217 |
| 20190612 | Spring | | | 1874 | |
| **Average** | | **3457** | **3111** | **1908** | **208** |

**Supplemental Table 4.** (this and next 2 pages) $\delta^{18}O$ and $\delta D$ measured in water samples from Alpena fountain, MIS Arena water and specific locations in the arena (when available), the Alcove groundwater source of the sinkhole, and the surface of the lake above the sinkhole. Alcove waters (45° 11.900'N, 83° 19.639'W) and Alpena fountain were sampled in the same locations. Surface waters were sampled above the sinkhole arena.

| Year | Date | Season | Sample | Location | Specific Site | $\delta^{18}O$(‰) | $\delta D$ (‰) |
|---|---|---|---|---|---|---|---|
| 2015 | 20151007 | Autumn | ALP.2015.008 | Alpena | | -12.2 | -83.8 |
| 2015 | 20151007 | Autumn | ALP.2015.014 | Alpena | | -12.1 | -82.6 |
| 2017 | 20170810 | Summer | ALP.2017.Aug | Alpena | | -12.4 | -86.9 |
| 2017 | 20170912 | Autumn | ALP.2017.Sept | Alpena | | -12.3 | -84.2 |
| 2018 | 20180710 | Summer | ALP.2018.July | Alpena | | -12.1 | -83.3 |
| 2015 | 20150604 | Spring | MIS.2015.008 | Alcove | | -11.4 | -78.0 |
| 2015 | 20150731 | Summer | MIS.2015.047 | Alcove | | -11.5 | -78.3 |
| 2015 | 20151007 | Autumn | MIS.2015.079 | Alcove | | -10.1 | -71.0 |
| 2016 | 20160609 | Spring | MIS2016.001 | Alcove | | -11.4 | -77.5 |
| 2016 | 20160928 | Autumn | MIS.2016.322 | Alcove | | -11.3 | -77.7 |
| 2017 | 20170531 | Spring | MIS.2017.005 | Alcove | | -11.5 | -78.3 |
| 2017 | 20170809 | Summer | MIS.2017.062 | Alcove | | -11.5 | -78.5 |
| 2017 | 20170810 | Summer | MIS.2017.080 | Alcove | | -11.6 | -79.1 |
| 2015 | 20150731 | Summer | MIS.2015.042 | Arena | | -9.7 | -68.8 |
| 2015 | 20151007 | Autumn | MIS.2015.077 | Arena | | -10.4 | -71.9 |
| 2016 | 20160609 | Spring | MIS2016.009 | Arena | 45° 11.918'N, 83° 19.662'W | -9.7 | -68.0 |
| 2016 | 20160609 | Spring | MIS2016.016 | Arena | 45° 11.944'N, 83° 19.689'W | -9.2 | -65.9 |

| | | | | | | | |
|---|---|---|---|---|---|---|---|
| 2016 | 20160609 | Spring | MIS2016.023 | Arena | 45° 11.949'N, 83° 19.649'W | -9.0 | -65.1 |
| 2016 | 20160928 | Autumn | MIS.2016.Arena D3-4 | Arena | 45° 11.912'N, 83° 19.661'W | -9.3 | -65.2 |
| 2016 | 20160928 | Autumn | MIS.2016.325 | Arena | 45° 11.925'N, 83° 19.649'W | -9.0 | -64.3 |
| 2016 | 20160928 | Autumn | MIS.2016.328 | Arena | 45° 11.915'N, 83° 19.654'W | -9.5 | -67.2 |
| 2017 | 20170531 | Spring | MIS.2017.012 | Arena | 45° 11.932'N, 83° 19.677'W | -8.8 | -63.0 |
| 2017 | 20170531 | Spring | MIS.2017.013 | Arena | 45° 11.932'N, 83° 19.677'W | -8.8 | -62.3 |
| 2017 | 20170810 | Summer | MIS.2017.083 | Arena | 45° 11.932'N, 83° 19.677'W | -10.3 | -71.4 |
| 2018 | 20180710 | Summer | P7.2018.July | Arena | 45° 11.932'N, 83° 19.677'W | -9.9 | -70.0 |
| 2015 | 20150604 | Spring | MIS.2015.001 | Surface | | -7.1 | -52.9 |
| 2015 | 20151007 | Autumn | MIS.2015.073 | Surface | | -7.1 | -53.2 |
| 2016 | 20160609 | Spring | MIS2016.026 | Surface | | -7.1 | -52.5 |
| 2016 | 20160928 | Autumn | MIS.2016.319 | Surface | | -7.0 | -52.0 |
| 2017 | 20170531 | Spring | MIS.2017.002 | Surface | | -7.1 | -52.4 |
| 2017 | 20170809 | Summer | MIS.2017.056 | Surface | | -7.2 | -52.4 |

| | | | | | | | |
|---|---|---|---|---|---|---|---|
| 2017 | 20170810 | Summer | MIS.2017.<br>077 | Surface | | -7.2 | -53.2 |
| 2018 | 20180710 | Summer | SFC.2018.<br>July | Surface | | -7.2 | -54.1 |

**Supplemental Table 5.** (this and next page) The average and standard deviation of relative abundances of key taxonomic groups by season unit.

| Taxon | Season | Mean (%) | Standard Deviation (%) |
|---|---|---|---|
| Cyanobacteria;*Phormidium* | Spring | 33.55 | 15.94 |
| Cyanobacteria;*Phormidium* | Summer | 35.55 | 23.92 |
| Cyanobacteria;*Phormidium* | Autumn | 4.19 | 3.96 |
| Cyanobacteria;*Planktothrix* | Spring | 8.05 | 7.83 |
| Cyanobacteria;*Planktothrix* | Summer | 1.88 | 4.36 |
| Cyanobacteria;*Planktothrix* | Autumn | 8.43 | 11.54 |
| Cyanobacteria;*Spirulina* | Spring | 0.02 | 0.03 |
| Cyanobacteria;*Spirulina* | Summer | 0.04 | 0.09 |
| Cyanobacteria;*Spirulina* | Autumn | 0.55 | 1.65 |
| Cyanobacteria;*Pseudanabaena* | Spring | 0.00 | 0.01 |
| Cyanobacteria;*Pseudanabaena* | Summer | 0.01 | 0.03 |
| Cyanobacteria;*Pseudanabaena* | Autumn | 0.15 | 0.30 |
| Other Cyanobacteria | Spring | 4.36 | 2.48 |
| Other Cyanobacteria | Summer | 2.47 | 1.96 |
| Other Cyanobacteria | Autumn | 2.43 | 1.42 |
| Epsilonproteobacteria S-oxidizers | Spring | 4.81 | 5.07 |
| Epsilonproteobacteria S-oxidizers | Summer | 2.41 | 1.93 |
| Epsilonproteobacteria S-oxidizers | Autumn | 3.11 | 1.66 |
| Other Epsilonproteobacteria | Spring | 0.06 | 0.08 |
| Other Epsilonproteobacteria | Summer | 0.22 | 0.30 |
| Other Epsilonproteobacteria | Autumn | 0.96 | 1.26 |
| Gammaproteobacteria;*Beggiatoa* | Spring | 4.78 | 7.40 |
| Gammaproteobacteria;*Beggiatoa* | Summer | 5.25 | 4.11 |
| Gammaproteobacteria;*Beggiatoa* | Autumn | 10.79 | 12.19 |
| Other Gammaproteobacteria | Spring | 2.27 | 1.46 |
| Other Gammaproteobacteria | Summer | 2.20 | 1.89 |
| Other Gammaproteobacteria | Autumn | 5.55 | 2.93 |
| Deltaproteobacteria;*Desulfonema* | Spring | 0.34 | 0.45 |
| Deltaproteobacteria;*Desulfonema* | Summer | 5.95 | 5.95 |
| Deltaproteobacteria;*Desulfonema* | Autumn | 6.06 | 4.97 |
| Deltaproteobacteria;*Desulfocapsa* | Spring | 2.67 | 1.99 |
| Deltaproteobacteria;*Desulfocapsa* | Summer | 3.00 | 2.57 |
| Deltaproteobacteria;*Desulfocapsa* | Autumn | 0.94 | 0.88 |
| Other Deltaproteobacteria | Spring | 1.85 | 1.19 |
| Other Deltaproteobacteria | Summer | 3.06 | 2.53 |
| Other Deltaproteobacteria | Autumn | 5.76 | 2.03 |
| Other Proteobacteria | Spring | 16.44 | 5.23 |
| Other Proteobacteria | Summer | 9.13 | 5.27 |
| Other Proteobacteria | Autumn | 12.25 | 6.19 |
| Acidobacteria | Spring | 0.33 | 0.17 |
| Acidobacteria | Summer | 0.52 | 0.55 |
| Acidobacteria | Autumn | 1.09 | 0.38 |
| Bacteroidetes | Spring | 14.73 | 4.72 |
| Bacteroidetes | Summer | 15.52 | 6.44 |
| Bacteroidetes | Autumn | 18.31 | 6.92 |
| Chlorobi | Spring | 0.13 | 0.08 |
| Chlorobi | Summer | 0.31 | 0.39 |
| Chlorobi | Autumn | 0.78 | 0.39 |

| | | | |
|---|---|---|---|
| Chloroflexi | Spring | 0.23 | 0.11 |
| Chloroflexi | Summer | 0.94 | 1.37 |
| Chloroflexi | Autumn | 1.75 | 0.68 |
| Firmicutes | Spring | 1.16 | 0.87 |
| Firmicutes | Summer | 3.46 | 5.56 |
| Firmicutes | Autumn | 2.80 | 1.50 |
| Planctomycetes | Spring | 0.24 | 0.15 |
| Planctomycetes | Summer | 0.93 | 1.00 |
| Planctomycetes | Autumn | 1.60 | 0.73 |
| Spirochaetes | Spring | 1.12 | 1.07 |
| Spirochaetes | Summer | 1.98 | 1.56 |
| Spirochaetes | Autumn | 2.87 | 1.46 |
| Verrucomicrobia | Spring | 1.42 | 0.96 |
| Verrucomicrobia | Summer | 2.07 | 2.66 |
| Verrucomicrobia | Autumn | 5.07 | 2.46 |

**Supplemental Table 6.** (this and next 2 pages) The average and standard deviation of relative abundances of Deltaproteobacterial putative sulfate reducers in each season unit.

| Taxon | Season | Mean (%) | Standard Deviation (%) |
|---|---|---|---|
| Desulfarculales;Desulfarculaceae;Desulfarculus | Spring | 0.00 | 0.00 |
| Desulfarculales;Desulfarculaceae;Desulfarculus | Summer | 0.00 | 0.00 |
| Desulfarculales;Desulfarculaceae;Desulfarculus | Autumn | 0.00 | 0.00 |
| Desulfarculales;Desulfarculaceae;Other | Spring | 0.00 | 0.01 |
| Desulfarculales;Desulfarculaceae;Other | Summer | 0.02 | 0.05 |
| Desulfarculales;Desulfarculaceae;Other | Autumn | 0.06 | 0.03 |
| Desulfobacterales;Desulfobacteraceae;Desulfatibacillum | Spring | 0.00 | 0.00 |
| Desulfobacterales;Desulfobacteraceae;Desulfatibacillum | Summer | 0.00 | 0.01 |
| Desulfobacterales;Desulfobacteraceae;Desulfatibacillum | Autumn | 0.00 | 0.00 |
| Desulfobacterales;Desulfobacteraceae;Desulfatiferula | Spring | 0.00 | 0.01 |
| Desulfobacterales;Desulfobacteraceae;Desulfatiferula | Summer | 0.01 | 0.02 |
| Desulfobacterales;Desulfobacteraceae;Desulfatiferula | Autumn | 0.03 | 0.02 |
| Desulfobacterales;Desulfobacteraceae;Desulfobacter | Spring | 0.01 | 0.02 |
| Desulfobacterales;Desulfobacteraceae;Desulfobacter | Summer | 0.00 | 0.01 |
| Desulfobacterales;Desulfobacteraceae;Desulfobacter | Autumn | 0.03 | 0.03 |
| Desulfobacterales;Desulfobacteraceae;Desulfobacterium | Spring | 0.19 | 0.15 |
| Desulfobacterales;Desulfobacteraceae;Desulfobacterium | Summer | 0.34 | 0.40 |
| Desulfobacterales;Desulfobacteraceae;Desulfobacterium | Autumn | 0.85 | 0.52 |
| Desulfobacterales;Desulfobacteraceae;Desulfobacula | Spring | 0.03 | 0.02 |
| Desulfobacterales;Desulfobacteraceae;Desulfobacula | Summer | 0.12 | 0.12 |
| Desulfobacterales;Desulfobacteraceae;Desulfobacula | Autumn | 0.31 | 0.21 |
| Desulfobacterales;Desulfobacteraceae;Desulfococcus | Spring | 0.00 | 0.00 |
| Desulfobacterales;Desulfobacteraceae;Desulfococcus | Summer | 0.00 | 0.00 |
| Desulfobacterales;Desulfobacteraceae;Desulfococcus | Autumn | 0.00 | 0.00 |
| Desulfobacterales;Desulfobacteraceae;Desulfofaba | Spring | 0.00 | 0.00 |
| Desulfobacterales;Desulfobacteraceae;Desulfofaba | Summer | 0.00 | 0.00 |
| Desulfobacterales;Desulfobacteraceae;Desulfofaba | Autumn | 0.00 | 0.00 |
| Desulfobacterales;Desulfobacteraceae;Desulfonema | Spring | 0.84 | 1.62 |
| Desulfobacterales;Desulfobacteraceae;Desulfonema | Summer | 5.29 | 5.57 |
| Desulfobacterales;Desulfobacteraceae;Desulfonema | Autumn | 6.06 | 4.97 |
| Desulfobacterales;Desulfobacteraceae;Desulforegula | Spring | 0.00 | 0.00 |
| Desulfobacterales;Desulfobacteraceae;Desulforegula | Summer | 0.00 | 0.00 |
| Desulfobacterales;Desulfobacteraceae;Desulforegula | Autumn | 0.00 | 0.01 |
| Desulfobacterales;Desulfobacteraceae;Desulfosarcina | Spring | 0.02 | 0.03 |
| Desulfobacterales;Desulfobacteraceae;Desulfosarcina | Summer | 0.07 | 0.19 |
| Desulfobacterales;Desulfobacteraceae;Desulfosarcina | Autumn | 0.15 | 0.18 |
| Desulfobacterales;Desulfobacteraceae;Desulfotignum | Spring | 0.00 | 0.00 |
| Desulfobacterales;Desulfobacteraceae;Desulfotignum | Summer | 0.00 | 0.01 |
| Desulfobacterales;Desulfobacteraceae;Desulfotignum | Autumn | 0.01 | 0.01 |
| Desulfobacterales;Desulfobacteraceae;Other | Spring | 0.03 | 0.03 |
| Desulfobacterales;Desulfobacteraceae;Other | Summer | 0.32 | 0.83 |
| Desulfobacterales;Desulfobacteraceae;Other | Autumn | 0.24 | 0.16 |
| Desulfobacterales;Desulfobulbaceae;Desulfobacterium | Spring | 0.22 | 0.22 |
| Desulfobacterales;Desulfobulbaceae;Desulfobacterium | Summer | 0.48 | 0.56 |
| Desulfobacterales;Desulfobulbaceae;Desulfobacterium | Autumn | 0.59 | 0.35 |
| Desulfobacterales;Desulfobulbaceae;Desulfobulbus | Spring | 0.14 | 0.20 |
| Desulfobacterales;Desulfobulbaceae;Desulfobulbus | Summer | 0.10 | 0.09 |
| Desulfobacterales;Desulfobulbaceae;Desulfobulbus | Autumn | 0.20 | 0.12 |

| | | | |
|---|---|---|---|
| Desulfobacterales;Desulfobulbaceae;Desulfocapsa | Spring | 2.40 | 1.84 |
| Desulfobacterales;Desulfobulbaceae;Desulfocapsa | Summer | 2.72 | 2.38 |
| Desulfobacterales;Desulfobulbaceae;Desulfocapsa | Autumn | 0.94 | 0.88 |
| Desulfobacterales;Desulfobulbaceae;Desulfopila | Spring | 0.00 | 0.00 |
| Desulfobacterales;Desulfobulbaceae;Desulfopila | Summer | 0.00 | 0.00 |
| Desulfobacterales;Desulfobulbaceae;Desulfopila | Autumn | 0.00 | 0.00 |
| Desulfobacterales;Desulfobulbaceae;Desulforhopalus | Spring | 0.01 | 0.02 |
| Desulfobacterales;Desulfobulbaceae;Desulforhopalus | Summer | 0.02 | 0.04 |
| Desulfobacterales;Desulfobulbaceae;Desulforhopalus | Autumn | 0.03 | 0.02 |
| Desulfobacterales;Desulfobulbaceae;Desulfotalea | Spring | 0.00 | 0.00 |
| Desulfobacterales;Desulfobulbaceae;Desulfotalea | Summer | 0.00 | 0.00 |
| Desulfobacterales;Desulfobulbaceae;Desulfotalea | Autumn | 0.00 | 0.00 |
| Desulfobacterales;Desulfobulbaceae;Desulfurivibrio | Spring | 0.00 | 0.00 |
| Desulfobacterales;Desulfobulbaceae;Desulfurivibrio | Summer | 0.00 | 0.00 |
| Desulfobacterales;Desulfobulbaceae;Desulfurivibrio | Autumn | 0.00 | 0.00 |
| Desulfobacterales;Desulfobulbaceae;Other | Spring | 0.00 | 0.00 |
| Desulfobacterales;Desulfobulbaceae;Other | Summer | 0.00 | 0.00 |
| Desulfobacterales;Desulfobulbaceae;Other | Autumn | 0.00 | 0.00 |
| Desulfobacterales;Nitrospinaceae;Nitrospinaceae | Spring | 0.00 | 0.00 |
| Desulfobacterales;Nitrospinaceae;Nitrospinaceae | Summer | 0.00 | 0.01 |
| Desulfobacterales;Nitrospinaceae;Nitrospinaceae | Autumn | 0.00 | 0.00 |
| Desulfobacterales;Other | Spring | 0.00 | 0.01 |
| Desulfobacterales;Other | Summer | 0.02 | 0.04 |
| Desulfobacterales;Other | Autumn | 0.06 | 0.06 |
| Desulfovibrionales;Desulfohalobiaceae;Desulfovermiculus | Spring | 0.00 | 0.00 |
| Desulfovibrionales;Desulfohalobiaceae;Desulfovermiculus | Summer | 0.01 | 0.01 |
| Desulfovibrionales;Desulfohalobiaceae;Desulfovermiculus | Autumn | 0.03 | 0.02 |
| Desulfovibrionales;Desulfomicrobiaceae;Desulfomicrobium | Spring | 0.21 | 0.20 |
| Desulfovibrionales;Desulfomicrobiaceae;Desulfomicrobium | Summer | 0.18 | 0.25 |
| Desulfovibrionales;Desulfomicrobiaceae;Desulfomicrobium | Autumn | 0.22 | 0.19 |
| Desulfovibrionales;Desulfonatronaceae;Desulfonatronum | Spring | 0.00 | 0.00 |
| Desulfovibrionales;Desulfonatronaceae;Desulfonatronum | Summer | 0.00 | 0.00 |
| Desulfovibrionales;Desulfonatronaceae;Desulfonatronum | Autumn | 0.00 | 0.00 |
| Desulfovibrionales;Desulfovibrionaceae;Desulfovibrio | Spring | 0.04 | 0.06 |
| Desulfovibrionales;Desulfovibrionaceae;Desulfovibrio | Summer | 0.04 | 0.03 |
| Desulfovibrionales;Desulfovibrionaceae;Desulfovibrio | Autumn | 0.10 | 0.05 |
| Desulfurellales;Desulfurellaceae;Other | Spring | 0.00 | 0.00 |
| Desulfurellales;Desulfurellaceae;Other | Summer | 0.00 | 0.00 |
| Desulfurellales;Desulfurellaceae;Other | Autumn | 0.00 | 0.00 |
| Desulfuromonadales;Desulfuromonadaceae;Desulfuromonas | Spring | 0.00 | 0.00 |
| Desulfuromonadales;Desulfuromonadaceae;Desulfuromonas | Summer | 0.00 | 0.00 |
| Desulfuromonadales;Desulfuromonadaceae;Desulfuromonas | Autumn | 0.00 | 0.00 |
| Desulfuromonadales;Desulfuromonadaceae;Other | Spring | 0.00 | 0.00 |
| Desulfuromonadales;Desulfuromonadaceae;Other | Summer | 0.00 | 0.00 |
| Desulfuromonadales;Desulfuromonadaceae;Other | Autumn | 0.00 | 0.00 |
| Desulfuromonadales;Desulfuromonadaceae;Pelobacter | Spring | 0.02 | 0.03 |
| Desulfuromonadales;Desulfuromonadaceae;Pelobacter | Summer | 0.02 | 0.03 |
| Desulfuromonadales;Desulfuromonadaceae;Pelobacter | Autumn | 0.03 | 0.05 |
| Desulfuromonadales;Geobacteraceae;Geobacter | Spring | 0.32 | 0.39 |
| Desulfuromonadales;Geobacteraceae;Geobacter | Summer | 0.11 | 0.12 |
| Desulfuromonadales;Geobacteraceae;Geobacter | Autumn | 0.34 | 0.49 |
| Desulfuromonadales;Geobacteraceae;Geothermobacter | Spring | 0.00 | 0.00 |

| | | | |
|---|---|---|---|
| Desulfuromonadales;Geobacteraceae;Geothermobacter | Summer | 0.00 | 0.00 |
| Desulfuromonadales;Geobacteraceae;Geothermobacter | Autumn | 0.00 | 0.00 |
| Desulfuromonadales;Other | Spring | 0.00 | 0.01 |
| Desulfuromonadales;Other | Summer | 0.02 | 0.03 |
| Desulfuromonadales;Other | Autumn | 0.05 | 0.04 |
| Syntrophobacterales;Syntrophaceae;Desulfobacca | Spring | 0.00 | 0.00 |
| Syntrophobacterales;Syntrophaceae;Desulfobacca | Summer | 0.00 | 0.01 |
| Syntrophobacterales;Syntrophaceae;Desulfobacca | Autumn | 0.01 | 0.01 |
| Syntrophobacterales;Syntrophaceae;Desulfomonile | Spring | 0.01 | 0.01 |
| Syntrophobacterales;Syntrophaceae;Desulfomonile | Summer | 0.02 | 0.03 |
| Syntrophobacterales;Syntrophaceae;Desulfomonile | Autumn | 0.03 | 0.03 |
| Syntrophobacterales;Syntrophobacteraceae;Desulfovirga | Spring | 0.00 | 0.00 |
| Syntrophobacterales;Syntrophobacteraceae;Desulfovirga | Summer | 0.00 | 0.01 |
| Syntrophobacterales;Syntrophobacteraceae;Desulfovirga | Autumn | 0.01 | 0.01 |

**Supplemental Table 7.** (this and next 2 pages) Weighted averages of $log_2$-normalized abundances of proteins across each season. $Log_2$-normalized abundances and spectral abundance of proteins in samples from respective season units were used to generate weighted averages and standard deviations (when a protein was observed in more than 1 sample in each season unit). Proteins found to be significantly differentially abundant between spring and summer are noted with "a", between spring and autumn with "b", and between summer and autumn with "c".

| Protein | Short name | System | Organism | Spring | Summer | Autumn | Significant relationships |
|---|---|---|---|---|---|---|---|
| 3300002024_MIS_11767773 | GroES | Central metabolism | Phormidium | 0.43 ± 0.29 | 0.92 ± 0.29 | -2.87 ± 0.2 | bc |
| 3300002024_MIS_11047972 | L7/L12 | Central metabolism | Phormidium | -1.02 ± 0.9 | -0.17 ± 0.14 | -2.25 ± 0.32 | c |
| 3300002027_MIS_100694095 | groEL | Central metabolism | Phormidium | -0.62 ± 0.42 | 0.35 ± 0.31 | -2.59 ± 0.19 | bc |
| 3300002027_MIS_101997102 | glnA | Nitrogen metabolism | Phormidium | -3.22 | 0.76 ± 0.67 | | |
| 3300002027_MIS_100173914 | apcA | Photosynthesis | Phormidium | -2.03 ± 0.55 | 0.21 ± 0.67 | -3 ± 0.38 | ac |
| 3300002027_MIS_100173913 | apcB | Photosynthesis | Phormidium | -0.62 ± 0.15 | 0.15 ± 0.46 | -2.61 ± 0.64 | c |
| 3300002027_MIS_101595453 | apcF | Photosynthesis | Phormidium | -0.62 ± 0.01 | 0.17 ± 0.3 | -1.53 | a |
| 3300002027_MIS_101997101 | apcF | Photosynthesis | Phormidium | -2.21 | -0.52 | | |
| 3300002027_MIS_100523012 | cpcA | Photosynthesis | Phormidium | -1.01 ± 0.17 | -0.47 ± 0.55 | -3.65 ± 0.38 | bc |
| 3300002027_MIS_100523013 | cpcB | Photosynthesis | Phormidium | -1.8 ± 0.2 | 0.33 ± 0.38 | -0.94 ± 1.31 | a |
| 3300002027_MIS_101905321 | cpcB | Photosynthesis | Phormidium | -2.89 ± 0.58 | 0.1 ± 0.13 | -1.43 ± 0.49 | a |
| 3300002027_MIS_100217001 | CpeA | Photosynthesis | Phormidium | -2.52 ± 0.55 | 0.05 ± 0.31 | -4.04 ± 2.25 | a |
| 3300002027_MIS_101906971 | CpeA | Photosynthesis | Phormidium | 1 ± 0.33 | -2.22 ± 1.4 | -0.5 ± 0.45 | b |
| 3300002027_MIS_101906972 | CpeB | Photosynthesis | Phormidium | 1.55 ± 0.45 | -1.87 ± 1.34 | -0.76 ± 0.53 | ab |
| 3300002024_MIS_10075351 | cpeE | Photosynthesis | Phormidium | -0.92 ± 0.27 | 1.29 ± 0.28 | -2.77 | a |
| 3300002027_MIS_101945691 | cpeE | Photosynthesis | Phormidium | -0.83 ± 0.04 | 0.41 ± 0.15 | -2.34 ± 0.29 | abc |
| 3300002026_MIS_100342443 | psaB | Photosynthesis | Phormidium | -0.22 ± 0.68 | -0.09 ± 0.18 | -1.44 ± 0.24 | c |
| 3300002027_MIS_100129452 | psaD | Photosynthesis | Phormidium | -0.41 ± 1.11 | 0.45 ± 0.32 | -2.89 ± 0.13 | c |
| 3300002024_MIS_11788891 | SOD | Photosynthesis | Phormidium | 0.14 ± 0.25 | 0.07 ± 0.29 | | |
| 3300002027_MIS_101999963 | trxA | Photosynthesis | Phormidium | 0.35 ± 0.45 | 0.2 ± 0.3 | -3.32 ± 0.48 | bc |
| 3300002027_MIS_101849361 | CnaB | Central metabolism | Phormidium | 3.56 ± 0.01 | 1.33 ± 0.26 | | a |
| 3300002027_MIS_100753761 | AtpD | Central metabolism | Planktothrix | -1.99 ± 0.89 | 0.41 ± 0.49 | -2.26 ± 0.15 | c |
| 3300002026_MIS_100162309 | apcA | Photosynthesis | Planktothrix | 1.54 ± 0.54 | 0.7 ± 0.34 | -1.69 ± 0.19 | bc |
| 3300002026_MIS_1001623010 | apcB | Photosynthesis | Planktothrix | 0.12 ± 0.36 | 0.03 ± 0.28 | -1.7 ± 0.15 | bc |
| 3300002026_MIS_1001623011 | apcC | Photosynthesis | Planktothrix | 0.42 ± 0.74 | | | |
| 3300002027_MIS_100155491 | cpcB | Photosynthesis | Planktothrix | 2.42 ± 0.84 | -2.58 ± 1.29 | -1.45 ± 0.39 | ab |

| | | | | | | | |
|---|---|---|---|---|---|---|---|
| 3300002027_MIS_100354662 | trxA | Photosynthesis | Planktothrix | 2.95 ± 0.47 | | -0.11 ± 0.18 | b |
| 3300002027_MIS_100969412 | n/a | Unknown | Planktothrix | 3.23 ± 0.3 | -1.87 ± 2.42 | -0.8 ± 0.27 | b |
| 3300002026_MIS_1000534011 | groEL | Central metabolism | Pseudanabaena | -1.94 ± 0.66 | 0.4 ± 0.34 | -3.65 ± 1.02 | a |
| 3300002026_MIS_1001557613 | CpeB | Photosynthesis | Pseudanabaena | -2.91 ± 1.16 | -0.49 ± 0.33 | -3.74 ± 0.12 | c |
| 3300002024_MIS_10041832 | psbD | Photosynthesis | Pseudanabaena | -0.24 ± 0.53 | -2.62 ± 0.39 | -1.3 ± 0.29 | ac |
| 3300002027_MIS_101689572 | cpcA | Photosynthesis | Spirulina | 0.75 ± 0.5 | -2.32 ± 0.95 | -0.66 ± 0.15 | a |
| 3300002027_MIS_101689573 | cpcB | Photosynthesis | Spirulina | 0.78 ± 1.52 | -5.28 ± 1.1 | -2.76 ± 0.54 | ac |
| 3300002026_MIS_100206493 | n/a | Unknown | Unknown cyanobacterium | 3.54 ± 0.51 | -0.81 | -0.43 ± 0.48 | b |
| 3300002024_MIS_10280361 | AtpA | Central metabolism | Putative bacillariophyte diatom | -0.01 ± 0.24 | -3.19 | -1.43 ± 0.41 | b |
| 3300002026_MIS_1000043450 | rbcL | Carbon metabolism | Putative bacillariophyte diatom | 0.21 ± 0.18 | -3.21 | -0.92 ± 0.11 | b |
| 3300002026_MIS_1000010375 | AtpD | Central metabolism | Putative bacillariophyte diatom | -0.01 ± 0.25 | -3.37 ± 0.49 | -1.37 ± 0.48 | ac |
| 3300002026_MIS_1000769711 | CpcG | Photosynthesis | Putative chloroplast genome | 0.71 ± 1.31 | -4.04 ± 0.65 | -2.16 ± 0.35 | ac |
| 3300002026_MIS_1000769710 | pecA | Photosynthesis | Putative chloroplast genome | 1.8 ± 1.1 | -3.63 ± 1.36 | -2.34 ± 0.29 | a |
| 3300002026_MIS_1000265839 | psaA | Photosynthesis | Putative chloroplast genome | -0.18 ± 0.06 | | -1.03 ± 0.55 | |
| 3300002026_MIS_1000265837 | psaF | Photosynthesis | Putative chloroplast genome | -0.09 ± 0.31 | -4.79 ± 1.25 | -1.58 ± 0.66 | ac |
| 3300002026_MIS_1000265832 | psaL | Photosynthesis | Putative chloroplast genome | 0.03 ± 0.36 | -4.31 | -0.01 ± 0.2 | |
| 3300002024_MIS_10189961 | AtpA | Central metabolism | Anaerolinea | -0.74 ± 0.13 | 0.17 ± 0.2 | -1.92 ± 0.25 | abc |
| 3300002026_MIS_100100153 | n/a | Unknown | Bacteria | -2.13 ± 0.4 | -0.11 ± 0.31 | -2.2 ± 1.95 | a |
| 3300002024_MIS_10898911 | GapA | Calvin-Benson-Bassham Cycle | Beggiatoa | | -1.87 ± 1.11 | 1.36 ± 0.21 | c |
| 3300002026_MIS_1003449713 | GroES | Central metabolism | Beggiatoa | | -0.38 | 0.16 | |
| 3300002026_MIS_1000443213 | L14 | Central metabolism | Beggiatoa | | | 0.47 ± 0.21 | |
| 3300002027_MIS_101719261 | OMP | Central metabolism | Beggiatoa | | -2.48 ± 1.39 | 1.38 ± 0.39 | c |
| 3300002026_MIS_100330993 | glnA | Nitrogen metabolism | Beggiatoa | | | -0.93 | |
| 3300002024_MIS_11106572 | DsrC | Sulfur oxidation | Beggiatoa | | | 0.42 | |
| 3300002027_MIS_100764955 | DsrC | Sulfur oxidation | Beggiatoa | | -2.67 ± 1.21 | -0.97 ± 1.18 | |
| 3300002024_MIS_11408701 | rdsrA | Sulfur oxidation | Beggiatoa | | | -0.12 | |
| 3300002027_MIS_101406783 | rdsrA | Sulfur oxidation | Beggiatoa | | -2.13 ± 1.26 | 0.37 ± 0.98 | |
| 3300002026_MIS_1000340919 | fusA, GFM, EFG | Translation | Beggiatoa | | | 0.39 | |
| 3300002027_MIS_101075071 | n/a | Unknown | Beggiatoa | | -3.53 ± 0.65 | 0.69 ± 0.44 | c |

| | | | | | | | |
|---|---|---|---|---|---|---|---|
| 3300002027_MIS_100225507 | aprB | Sulfate/sulfite reduction | Desulfobacteraceae | | -0.06 ± 0.95 | -1.32 ± 0.18 | |
| 3300002024_MIS_11625411 | AtpD | Central metabolism | Desulfotalea | -3.75 | -2.25 ± 0.92 | 0.24 ± 0.48 | c |
| 3300002024_MIS_10135141 | dsrA | Sulfate/sulfite reduction | SRB | | -0.14 | | |
| 3300002027_MIS_101924972 | DsrC | Sulfate/sulfite reduction | SRB | | -0.08 ± 1.58 | | |
| 3300002026_MIS_100111357 | OMP | Central metabolism | Methylococcaceae | | -2.29 ± 0.85 | 1.93 ± 0.3 | c |
| 3300002024_MIS_10200811 | Fba | Carbon fixation | Rhodoferax | -2.7 ± 0.31 | 0.18 ± 0.5 | | a |
| 3300002027_MIS_100411962 | UBC | Central metabolism | Eukaryote | 0.01 | -3.64 ± 0.85 | 0 ± 0.3 | c |
| 3300002024_MIS_10574951 | ACTB_G1 | Signal transduction | Eukaryote | 1.09 ± 0.42 | -1.77 ± 0.3 | -1.49 ± 0.23 | ab |
| 3300002024_MIS_11767773 | GroES | Central metabolism | Phormidium | 0.43 ± 0.29 | 0.92 ± 0.29 | -2.87 ± 0.2 | bc |
| 3300002024_MIS_11047972 | L7/L12 | Central metabolism | Phormidium | -1.02 ± 0.9 | -0.17 ± 0.14 | -2.25 ± 0.32 | c |
| 3300002027_MIS_100694095 | groEL | Central metabolism | Phormidium | -0.62 ± 0.42 | 0.35 ± 0.31 | -2.59 ± 0.19 | bc |
| 3300002027_MIS_101997102 | glnA | Nitrogen metabolism | Phormidium | -3.22 | 0.76 ± 0.67 | | |
